# Enhanced thermal stability enables human mismatch-specific Thymine-DNA glycosylase to catalyse futile DNA repair

**DOI:** 10.1101/2024.05.21.595234

**Authors:** Diana Manapkyzy, Botagoz Joldybayeva, Alexander A. Ishchenko, Bakhyt T. Matkarimov, Dmitry O. Zharkov, Sabira Taipakova, Murat K. Saparbaev

## Abstract

Human thymine-DNA glycosylase (TDG) excises T mispaired with G in a CpG context to initiate the base excision repair (BER) pathway. TDG is also involved in epigenetic regulation of gene expression by participating in active DNA demethylation. Here we demonstrate that under extended incubation time the full-length TDG (TDG^FL^), but not its truncated catalytic domain TDG (TDG^cat^) or methyl-binding protein 4 (MBD4) DNA glycosylase, exhibits a significant excision activity towards T and C in regular non-damaged DNA duplex in TpG/CpA and CpG/CpG contexts. Time course of the cleavage product generation under single-turnover conditions shows that the maximal rate of base excision (*k*obs) of TDG^FL^-catalysed excision of T in the T•A base pair (0.0014 – 0.0069 min^-1^) is 85 - 330-fold lower than that of T in T•G mismatch (0.47 – 0.61 min^-1^). Unexpectedly, TDG^FL^, but not TDG^cat^, exhibits prolonged enzyme stability under 37°C when incubated in the presence of equimolar concentration of non-specific DNA duplex, suggesting that the disordered N- and C-terminal domains of TDG can interact with DNA and stabilize the overall conformation of the protein. Noteworthy, TDG^FL^ is able to excise 5-hydroxymethylcytosine (5hmC), but not 5-methylcytosine residues, from duplex DNA with the efficiency that could be physiologically relevant in post-mitotic cells. Our findings demonstrate that, under the experimental conditions used, TDG catalyses sequence context-dependent removal of T, C and 5hmC residues in regular DNA duplexes. We propose that *in vivo* this futile DNA repair may lead to formation of persistent single-strand breaks in non-methylated regions of chromosomal DNA.

## Introduction

Post-replicative methylation of cytosine at the C5 position (5mC) together with its erasure are essential epigenetic processes in the course of organism development, cell differentiation, genomic imprinting and suppression of mobile elements [1–3]. A major drawback of DNA methylation is that spontaneous deamination of 5mC generates thymine, resulting in a T•G mismatch. If not repaired, these stray thymines result in C→T transition mutations at CpG dinucleotides. It is thought that the depleted CpG content in mammalian genomes is due to this high intrinsic mutability of 5mC [4]. In mammalian cells, both the methyl-binding domain protein 4 (MBD4/MED1) and mismatch-specific thymine-DNA glycosylase (TDG) counteract the mutagenic impact of 5mC deamination by excising thymine from T•G mispairs in a CpG context, which is then replaced with regular cytosine completing the base excision repair (BER) pathway [5, 6]. In BER, a DNA glycosylase recognizes the abnormal base and cleaves the glycosidic bond, generating an abasic site, which in turn is repaired by an apurinic/apyrimidinic (AP) endonuclease [7, 8]. Human TDG and MBD4 were first biochemically characterized for their ability to remove T mispaired with G. A more detailed characterization showed that TDG exhibits a wide DNA substrate specificity, excising 3,*N*^4^-ethenocytosine (εC) [9, 10], thymine glycol [11], 5-hydroxycytosine [12], 7,8-dihydro-8-oxoadenine (8oxoA) [13], mismatched uracil [14] and its derivatives modified at C5 [15]. In contrast, MBD4 has a narrow DNA substrate specificity, excising, beyond T, only uracil, 5-fluorouracil and 5-hydroxymethyluracil opposite guanine in duplex DNA [6, 16, 17]. Importantly, TDG is highly conserved in vertebrates.

In mammalian cells, TDG is tightly associated with euchromatin and binds CpG-rich promoters of transcribed genes to scan CpG sites for mismatches and to protect CpG from *de novo* DNA methylation [18, 19]. Studies of the mechanisms of active DNA demethylation in mammals have demonstrated that the Ten-eleven translocation family proteins (TETs) oxidize 5mC and the resulting products are removed by TDG. TET dioxygenases convert 5mC to 5-hydroxymethylcytosine (5hmC) and then to 5-formylcytosine (5fC) and 5-carboxylcytosine (5caC) both *in vitro* and *in vivo* [20–23]. TDG efficiently excises 5fC and 5caC nucleobases, but not 5hmC, in a CpG context, suggesting direct involvement of the TDG-initiated BER in the active erasure of 5mC from the genome [22, 24]. Furthermore, TDG knockout in mice is embryonic lethal due to the aberrant *de novo* DNA methylation of CpG islands (CGIs) in promoters of developmental genes, which results in a failure to establish and maintain cell type-specific gene expression programs during embryonic development [18, 19]. Noteworthy, ectopic overexpression of TDG hinders S-phase progression and cell proliferation, since this protein is strictly cell-cycle regulated: it is present in cells throughout the G2-M and G1 phases, but rapidly disappears in the S phase [25].

TDG consists of a conserved, well-folded catalytic core and extended, likely disordered N- and C-terminal tails encompassing about a half of the protein’s total length. This organization makes TDG quite a labile protein and notably affects its activity and specificity under different conditions. Studies in Drohat’s laboratory revealed that TDG is unstable at 37°C and prone to irreversible unfolding coupled to protein aggregation; whereas at 22°C the protein remains stable for at least 5 h [26]. On the other hand, G/T specific activity of TDG depends strongly on the temperature with an 11-fold increase from 5°C to 37°C, as compared to uracil-DNA glycosylase activity, which increases only threefold [26]. Intriguingly, the presence of a 10-fold molar excess of non-specific DNA in reaction buffer stabilizes TDG conformation against heat-induced unfolding for over 2 h at 37°C, suggesting that DNA binding prevents protein aggregation. Based on these observations, Drohat’s laboratory extensively characterized the DNA substrate specificity of TDG under room temperature conditions. Importantly, they studied TDG-catalysed aberrant removal of T paired with A, as well as C, 5mC and 5hmC residues paired with G in non-damaged DNA duplexes, and observed no significant cleavage after 1-2 h incubation at 37°C and even after 18 h or longer incubation at 22°C [12, 24]. For example, after incubation for 50 h at 22°C, TDG was able to excise T opposite to A in TpG/CpA context, but with extremely low efficiency, ∼10^4.3^-fold lower than that of T in G•T base pair [27, 28].

5hmC, now often regarded as a “6^th^ base” of mammalian DNA, is found at the steady-state level several orders of magnitude higher than 5fC and 5caC [29, 30]. Removal of TET-generated 5fC and 5caC by TDG and their replacement with regular cytosine via the BER pathway is required for active DNA demethylation and cell reprogramming during embryonic development. Yet the generation of 5hmC itself appears insufficient to drive rapid expression changes, or even induce significant DNA demethylation when reprogramming genomic loci and establishing cell lineages [31]. Despite being an intermediate product, 5hmC persist in many cell types and remains stable during cell divisions [32, 33]. Conversion of 5mC to 5hmC by TET enzymes occurs slowly during the first 30 h after DNA replication [32], and because of this, the global genomic level of 5hmC depends on the rate of cell proliferation, being the highest in the differentiated non-proliferating cell types and lowest in all studied cancers [34, 35]. 5hmC residues are typically enriched at enhancers and gene bodies of transcriptionally active and tissue-specific genes and appears to be a stable epigenetic mark that plays important role in the context of cell differentiation [36–39]. At present, the mechanistic aspects of 5hmC engagement in epigenetic regulation of gene expression remain murky. It is thought that 5hmC is not repaired by known DNA glycosylases.

Although the major role of DNA repair is to protect cells from DNA damage and prevent mutations, evidence have accumulated showing that mishandling of certain DNA lesions or even normal DNA could lead to faulty DNA repair and contribute to age-related disorders such as cancer and neurodegenerative diseases. For example, human alkylpurine DNA glycosylase (ANPG) can initiate futile BER by removing regular purines from non-damaged DNA [40], and increased level of ANPG is associated with increased risk of lung cancer [41]. Furthermore, previously we had shown that TDG and MBD4 catalyse aberrant excision of T paired with damaged adenine residues [42]. TDG targets the non-damaged DNA strand and efficiently excises T opposite to hypoxanthine (Hx), 1,*N*^6^-ethenoadenine (εA), 8-oxoadenine (8oxoA) and AP site in the TpG/CpX sequence context, where X is a modified residue. MBD4 removes T only from pairs with εA, but not with Hx or other adenine modifications. *In vitro* reconstitution showed that TDG-catalysed aberrant excision of a regular thymine opposite to Hx initiates repair synthesis that uses the damaged DNA template, which in turn leads to TpG→CpG and CpA→CpG mutations with no need for DNA replication [42]. These observations suggest that the DNA repair machinery can target the non-damaged DNA strand in an aberrant manner and promote genome instability in the presence of unrepaired DNA lesions.

Regulation of transcription via DNA methylation and demethylation is a common form of gene control in eukaryotes. The activity of transcription factors dependent on the methylation status allows genes to be specifically regulated via erasure and re-establishment of 5mC marks during development and cell differentiation. Enhancers are *cis*-acting regulatory DNA sequences (50-1500 bp) that, when bound by specific transcription factors, boost the transcription of their functionally associated genes. Physically, they may be located far away from the transcription start site and regulate the expression by DNA looping to interact with the target promoters. Noteworthy, the regulatory function of enhancers can be limited to a particular cell type, time point in the development, or physiological or environmental conditions. It was proposed that the generation of transient DNA strand breaks in the regulatory regions of genes can activate promoter and enhancer elements [43]. Recent studies demonstrated that post-mitotic neurons accumulate high levels of persistent single-strand DNA breaks (SSBs) in the enhancers at CpG dinucleotides associated with DNA demethylation, which control the expression of genes involved in cell identity [44, 45]. It has been further demonstrated that TDG/TET-mediated removal of 5mC residues is the source of these persistent SSBs in differentiated neurons and macrophages, suggesting that cycles of DNA methylation/demethylation are a major source of endogenous DNA damage in post-mitotic cells [46].

In the present work, during characterization of TDG-catalysed activities towards thymine paired with damaged adenine in duplex DNA, we found that upon long incubation at 37°C TDG can also excise regular pyrimidine residues in non-damaged DNA duplex. We characterise this futile DNA repair activity and provide biochemical evidences for a putative role of TDG in the generation of persistent single-strand DNA breaks and repair of 5hmC residues in post-mitotic cells.

## Material and Methods

### Proteins

Phage T4 polynucleotide kinase and and *E. coli* exonuclease III (Xth) were purchased from New England Biolabs (Evry, France). The *E. coli* BL21 (DE3) cells were purchased from Novagen-EMD4Biosciences (Merck Chemicals, Nottingham, UK). The purified BER proteins including truncated version of human uracil-DNA glycosylase (hUNGΔ84), *E. coli* endonuclease IV (Nfo) and human AP endonuclease 1 (APE1) were from laboratory stocks [47]. Expression and purification of TDG and MBD4 is described below. The activities of DNA glycosylases were verified using their canonical substrates and were tested just prior to use.

### Oligonucleotides

Sequences of the DNA and RNA oligonucleotides used in this work are shown in Table 1. All oligonucleotides containing modified bases and their complementary strands were purchased from Eurogentec (Seraing, Belgium). The oligonucleotides were 5’-end labelled with [γ-^32^P]-ATP (PerkinElmer, France) or labelled with fluorescent dye Cy5 (Cyanine5, far-red-fluorescent dye conjugated to the 5ʹ end of oligonucleotides) and then annealed with corresponding complementary strands as described previously [48]. The resulting oligonucleotide duplexes are referred to as X*•N, where X is a residue in the [^32^P]-labelled strand and N is a regular DNA base opposite to X in the complementary non-labelled strand.

**Table 1.**
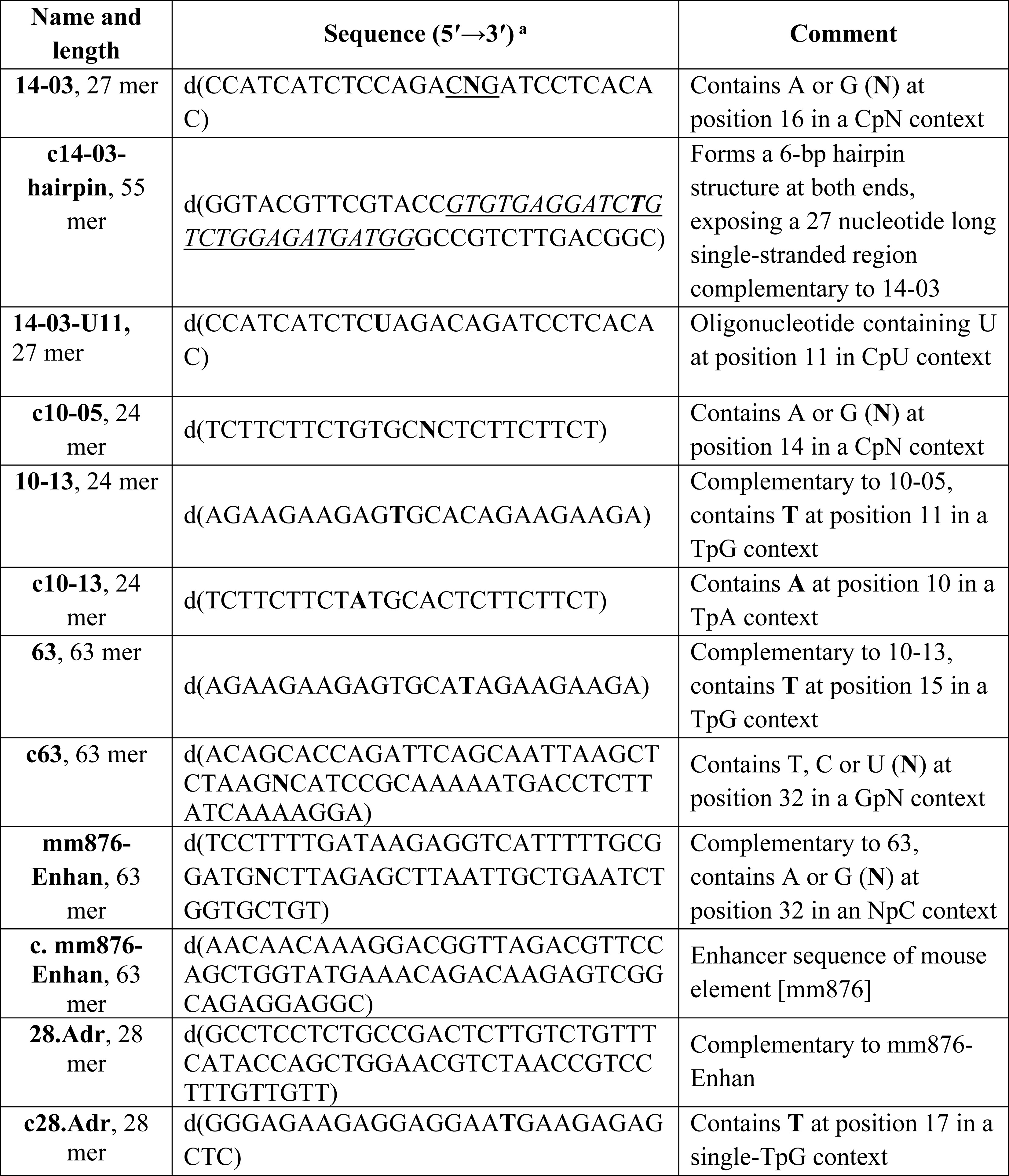
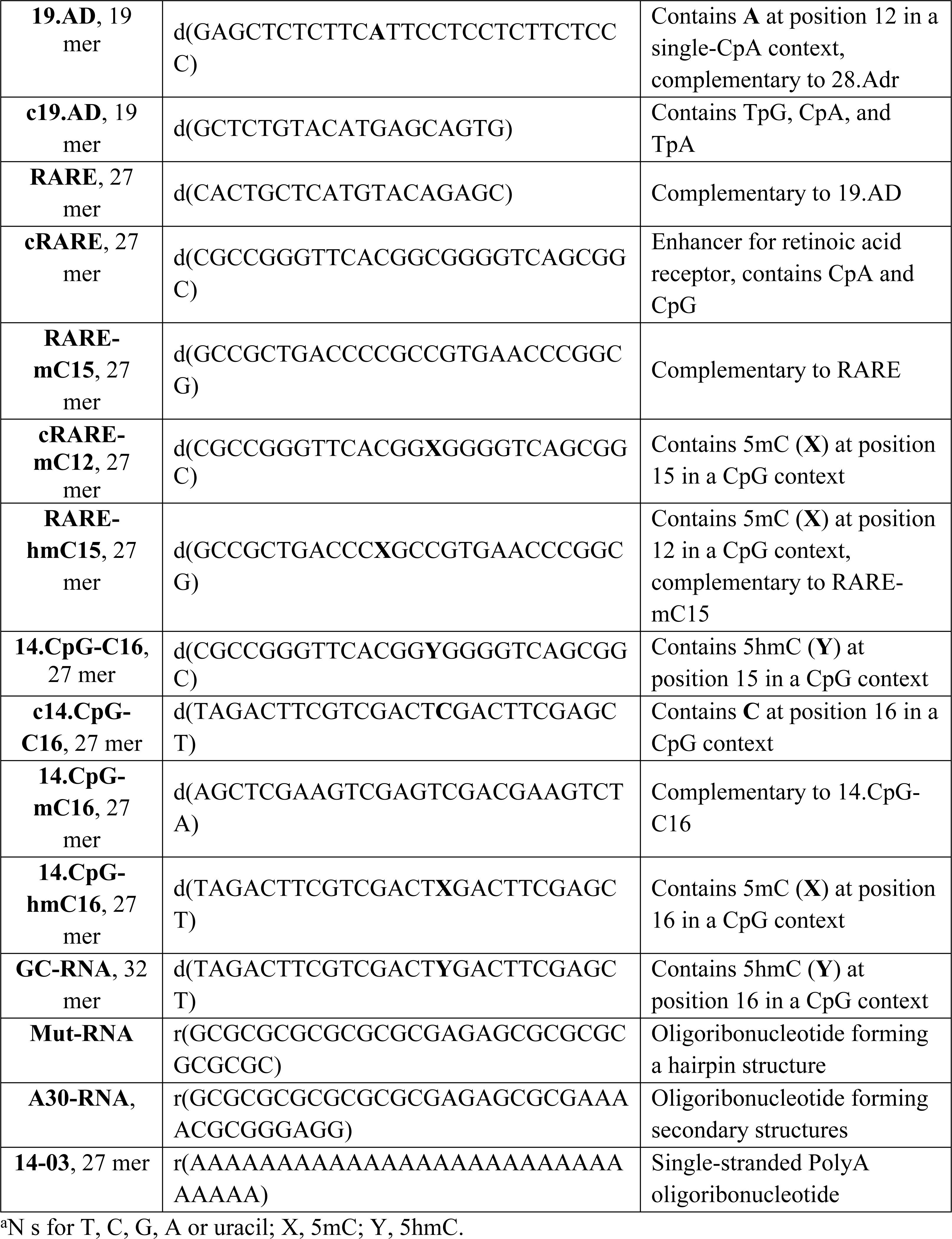
DNA sequence of the oligonucleotides used in the study.

### Expression and purification of TDG and MBD4

The expression and purification of TDG, TDG^cat^ and MBD4 proteins were produced as described previously [13, 47]. Briefly, Arctic Express (DE3) cells were transformed with the expression vectors pET28c-TDG (for full-length TDG protein), pET28c-TDG^cat^ (for catalytic domain TDG protein) and pET6H-MBD4 (for full-length MBD4 protein) and then grown on an orbital shaker at 37°C in the LB medium, supplemented with 50 µg•ml^-1^ of kanamycin or ampicillin, on an orbital shaker to OD600 nm = 0.6-0.8. Then the temperature was reduced to 12°C and the TDG^FL^, MBD4 and TDG^cat^ proteins expression was induced by 0.2 mM isopropyl β-D-1-thiogalactopyranoside (IPTG) and the cells were further grown for 15 h. The bacteria were harvested by centrifugation and the cell pellets were lysed using a French press at 18,000 psi in buffer containing 20 mM HEPES-KOH (pH 7.6), 40 mM NaCl and 0.025% NP-40, supplemented with Complete^TM^ Protease Inhibitor Cocktail (Roche Diagnostics, Switzerland). The lysates were cleared by centrifugation at 40,000 x *g* for 1 h at 4°C, the resulting supernatant was adjusted to 500 mM NaCl and 20 mM imidazole and loaded onto a HiTrap Chelating HP column (Amersham Biosciences, GE Healthcare). All purification procedures were carried out at 4°C. The column was washed with Buffer A (20 mM HEPES, 500 mM NaCl, 20 mM imidazole) and the bound proteins were eluted with a linear gradient of 20-500 mM imidazole in buffer A.

The proteins were further purified using Heparin HP affinity column. After HiTrap Chelating HP column fractions containing human protein were pooled and diluted ten-fold in buffer B (20 mM HEPES (pH 7.6), 50 mM NaCl and 0,0125% NP-40) and loaded to a 1 mL HiTrap-HeparinTM column (Amersham Biosciences, Orsay, France). Proteins bound to the column were eluted by a 50 to 800 mM NaCl gradient in Buffer B. The eluted fractions were analyzed by SDS-PAGE and the fractions containing the pure His-tagged TDG, TDG^cat^ and MBD4 proteins were stored at -80°C in 50% glycerol. The concentration of purified proteins was determined by the Bradford assay.

### DNA repair activity

Unless indicated otherwise, the standard reaction mixture (20 µl) for DNA repair assays contained 20 nM 5′-^32^P-labelled oligonucleotide duplex, 20 mM Tris–HCl (pH 8.0), 1 mM EDTA, 1 mM DTT, 100 µg•ml^−1^ BSA and 200 nM TDG^FL^, TDG^cat^ or MBD4. The reaction mixtures were incubated at 37°C for 5 to 60 min when measuring TDG-specific activities (T•G and U•G duplexes in which T- and U-containing strands were ^32^P-labelled) or 1 h and 18 h when measuring TDG-catalysed futile activity (T*•A, T•A*, C*•G and C•G* duplexes; the asterisk denotes the [^32^P]-labelled strand containing the specified base in a defined position). The reaction mixture for *E. coli* Nfo contained 50 mM KCl, 20 mM HEPES–KOH (pH 7.6), 100 µg•ml^−1^ BSA, 1 mM DTT and 1 nM Nfo. The reaction for *E. coli* exonuclease III (Xth) was performed in NEBuffer 1 (New England Biolabs) with 1 nM enzyme. The reaction mixture for the human uracil–DNA glycosylase (hUNGΔ84) contained 20 mM Tris–HCl (pH 8.1), 100 mM NaCl, 1 mM EDTA, 1 mM DTT, 100 µg•ml^−1^ BSA, and was incubated for 30 min at 37°C. All reactions were stopped by adding 10 µl of a stop solution consisting of 0.5% SDS and 20 mM EDTA. After incubation, the samples were treated either with 0.1 M NaOH for 3 min at 95°C and then neutralized by 0.1 M HCl, or with light piperidine (10% v/v piperidine at 37°C for 30 min) in order to cleave AP sites left after base excision by DNA glycosylases. To analyse the reaction products, the samples were desalted using Sephadex G25 DNA grade SF column (Amersham Biosciences) equilibrated in 7.5 M urea. The desalted reaction products were separated by electrophoresis in denaturing 20% (w/v) polyacrylamide gels (7 M urea, 0.5×TBE, 42°C). The gels were exposed to a Fuji FLA-3000 Phosphor Screen, then scanned with Typhoon FLA 9500 and quantified using Image Gauge V4.0 software. At least three independent experiments were conducted for all kinetic measurements. The release of T and C residues were measured by the cleavage of regular oligonucleotides containing T•A and C•G pairs, or oligonucleotides containing a single T•G or U•G mispair.

To generate size markers 20 nM [^32^P]-labelled oligonucleotide duplexes were incubated either with 10 nM of Nfo for 5-60 min at 37°C, or with 1 unit of Xth for 10 min at room T°C. The reactions were stopped by adding stop solution (0.1 M NaOH, 10mM EDTA, 0.25 % SDS) and heating at 95°C for 5 min. The size markers were also generated by using Fast Bisulfite Conversion Kit (Qiagen, Düsseldorf, Germany) and the hUNG protein. For this, 20 nM [^32^P]-labelled single-stranded oligonucleotides was treated with bisulfite according to manufacturer’s instructions to convert cytosine residues to uracil. After bisulfite treatment, the oligonucleotides were incubated with 10 nM hUNG at 37°C for 10 min to excise uracil residues followed by hot alkaline to cleave at AP sites.

### Single turnover kinetics

To obtain rate constants (*k*obs) that are not affected by enzyme-substrate association or by product inhibition, such that *k*obs reflects the maximal base excision rate (*k*obs ≈ *k*max), we used single-turnover kinetics under a large excess of the enzyme over the substrate ([E] >> [S] > *K*d). The time courses were performed in a large volume with 200 nM TDG and 20 nM substrate at 37°C. At each time point, a 20 µL aliquot was withdrawn and treated and analysed as described above. The data were fitted by non-linear regression to one-phase exponential association Equation S1 using GraphPad Prism 5 software.

[Fraction product] = *A*(1-exp(-*k*obs*t*)) (Eq. S1)

where *A* is the amplitude, *k*obs is the rate constant, and *t* is the reaction time (in minutes).

## Results

### Human native TDG excises regular pyrimidines from non-damaged duplex DNA and generates abasic sites

In our unpublished studies, we observed weak activity of native TDG towards regular DNA oligonucleotide duplexes upon long incubation time at 37°C, which we believed was due to contaminating non-specific nucleases. Here, we decided to examine whether TDG^FL^ can act on regular non-damaged DNA substrates, using catalytically inactive TDG^FL^-N140A mutant as a control for the presence of a contaminating activity from the overproducing cells. To avoid degradation of a DNA substrate by a putative nuclease activity, we constructed a 41-mer dumbbell-shaped duplex (dmbDNA) composed of a long 55-mer strand forming a hairpin structure at both ends and a complementary 27-mer oligonucleotide annealing in the middle (Fig. 1 and Table 1). The 27-mer fragment, which contained either A or G at position 16 or U at position 11, was hybridized to the 5′-^32^P-labelled 55-mer containing a regular T at position 26 opposite to A or G, to obtain the 41-mer dmbDNA referred as 14-03 in which the A residue was located in a CpA sequence context (Fig. 1B). To examine DNA glycosylase activity on both strands of a duplex, we also used dmbDNA with the 5′-^32^P-labelled 27-mer strand was hybridized with the non-labelled 55-mer hairpin oligonucleotide. The resulting dmbDNA substrates G16•T*, T•A*, A•T* and T•U11* (asterisk denotes the labelled DNA strand with the residue of interest) were incubated in the presence of TDG^FL^ or TDG^FL^-N140A for 1 h or 18 h at 37°C, followed by either light piperidine or hot alkaline treatment to cleave AP site generated by the DNA glycosylase action.

**Figure 1.**
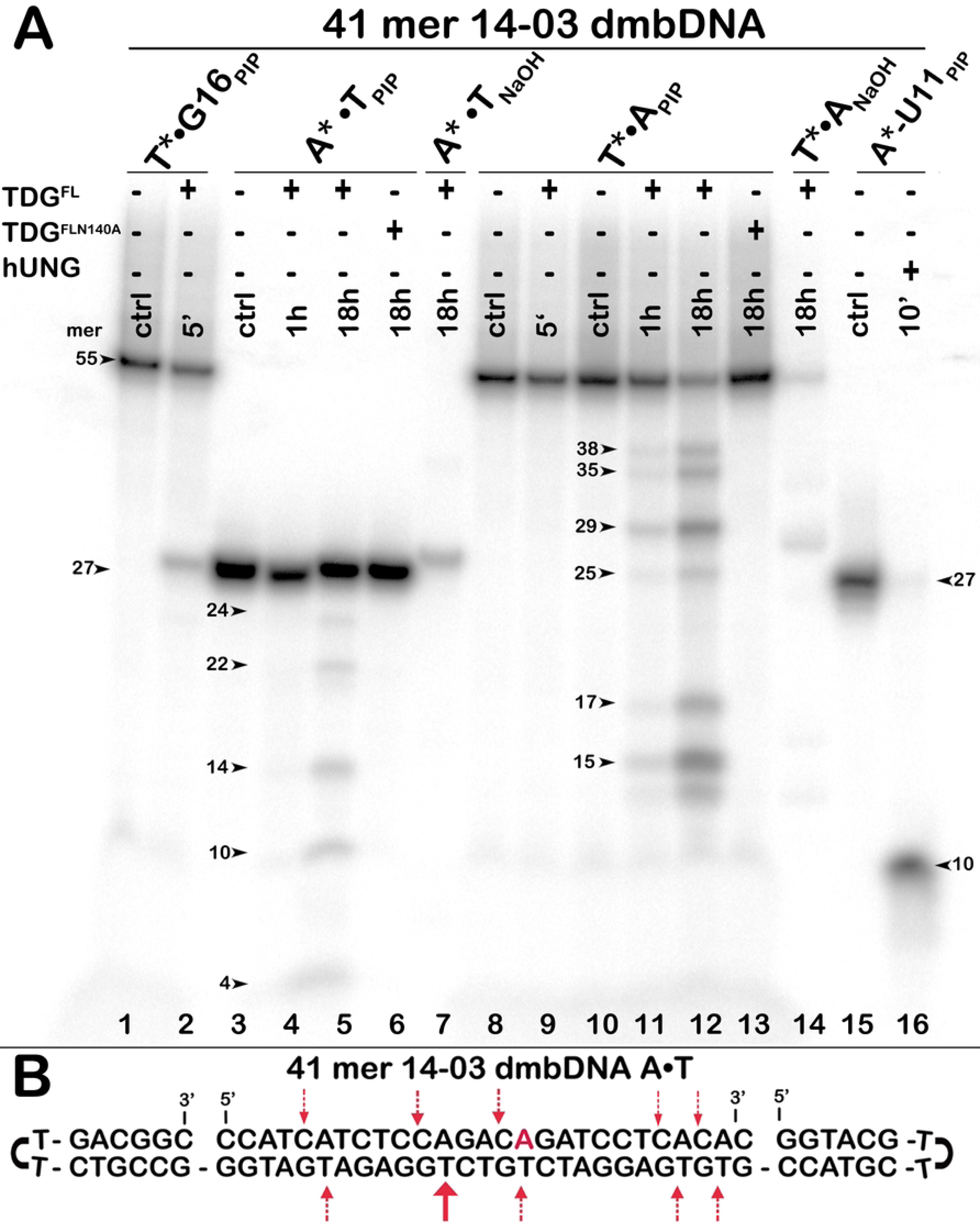
Action of native TDG^FL^ and catalytically dead mutant TDG^FL^-N140A on regular non-damaged 41 mer 14-03 A*•T and A•T* dmbDNA substrate, in which either top or bottom DNA strand is 5’-[^32^P]-labelled. (A) Denaturing PAGE analysis of cleavage products. Lane 1, control 41 mer G•T* dmbDNA containing T at position 26; lane 2, same as 1, but with TDG^FL^; lane 3, control 41 mer A*•T dmbDNA; lanes 4-5, same as 3, but with TDG^FL^; lane 6, same as 3, but with TDG^FL^-N140A; lane 7, same as 5, but treated with NaOH; lanes 8-14, 41 mer A•T* dmbDNA with 55 mer bottom strand labelled; lane 14, same as 12, but treated with NaOH; lane 15, 5’-[^32^P]-labeled 27 mer top strand containing U at position 11; lane 16, same as 15, but with hUNG. (B) Schematic representation of dmbDNA sequence with arrows depicting pyrimidines excised by TDG^FL^. For details, see Materials and Methods.

As shown in Figure 1, 5-min incubation of 14-03 G16•T* duplex with limited amount of TDG^FL^ resulted in the excision of T opposite to G and the generation of a 25-mer cleavage product (Fig. 1A, lane 2). To generate a 10-mer size marker we treated a 27-mer single-stranded oligonucleotide containing U at position 11 with hUNG (Fig. 1A, lane 16). Note that the 25-mer cleavage product (Fig. 1A, lane 2) migrates slower than the 27-mer oligonucleotide of the A*•T duplex and the 27-mer T•U11* duplex containing U (Fig. 1A, lanes 3 and 15), possibly due to the presence of a hairpin structure in the former. Unexpectedly, long 18-h incubation of TDG^FL^ with A*•T dmbDNA, in which the 27-mer strand with A at position 16 is labelled generates 24-, 22-, 14-, 10- and 4-mer cleavage fragments (Fig. 1A, lane 5), suggesting futile excision of regular cytosine residues opposite to G in a CpA context (Fig. 1B). Moreover, incubation of TDG^FL^ with A•T* dmbDNA, in which the 55-mer strand with T at position 26 is labelled, for 1 h and 18 h generated 38-, 35-, 29-, 25-, 17- and 15-mer cleavage fragments of varying intensities (Fig. 1A, lanes 11 and 12), implying the excision of regular T residues opposite to A at positions 39, 36, 30, 26, 18 and 16 in the hairpin oligonucleotide (Fig. 1B). Noteworthy, TDG^FL^ preferentially excised T from positions 30 and 16 of the 55-mer. Note that two control samples were treated by hot alkali instead of light piperidine, which resulted in a partial loss of radioactivity (Fig. 1A, lanes 7 and 14). Mutant TDG^FL^-N140A cleaved neither A*•T nor A•T* dmbDNA after 18-h incubation at 37°C (Fig. 1A, lanes 6 and 13), indicating that the futile excision of pyrimidines is not of the bacterial host origin, but is an intrinsic property of the human DNA glycosylase.

To further inquire into the nature of the observed enzymatic activity, we examined whether both native TDG^FL^ and its truncated version TDG^cat^ (residues 111–308) generate abasic sites after excision of regular pyrimidines. For this end, the products of reaction were analysed by denaturing PAGE before and after hot alkaline treatment. As shown in Figure 2, in the absence of hot alkaline treatment, incubation of TDG^FL^ with 14-03 A•T* dmbDNA for 18 h produced faint smeared cleavage fragments (Fig. 2, lane 7), suggesting that the enzyme cannot directly cleave the DNA duplex. As expected, when TDG^FL^ reaction products were treated by hot alkali, distinct cleavage fragments appeared (Fig. 2, lanes 8 and 9), suggesting that this pyrimidine-specific futile activity is not due to an AP lyase or DNA nuclease, but results from TDG^FL^ excising regular pyrimidines to generate AP sites in duplex DNA. Notably, TDG^cat^ exhibited dramatically lower futile activity towards 14-03 A•T* dmbDNA substrate as compared to its full-length counterpart (Fig. 2, lane 5 *vs* lane 9). After 18 h, TDG^cat^ excised only 0.8% of T at position 30 in the 55-mer strand of 14-03 A•T* dmbDNA, thus exhibiting 37-fold lower efficiency as compared to TDG^FL^, which excised 30.1% of T at the same position. These results suggest a possible role of non-catalytic N-terminal (residues 1–110) and C-terminal (residues 309–410) tails of human TDG in the futile DNA repair.

**Figure 2.**
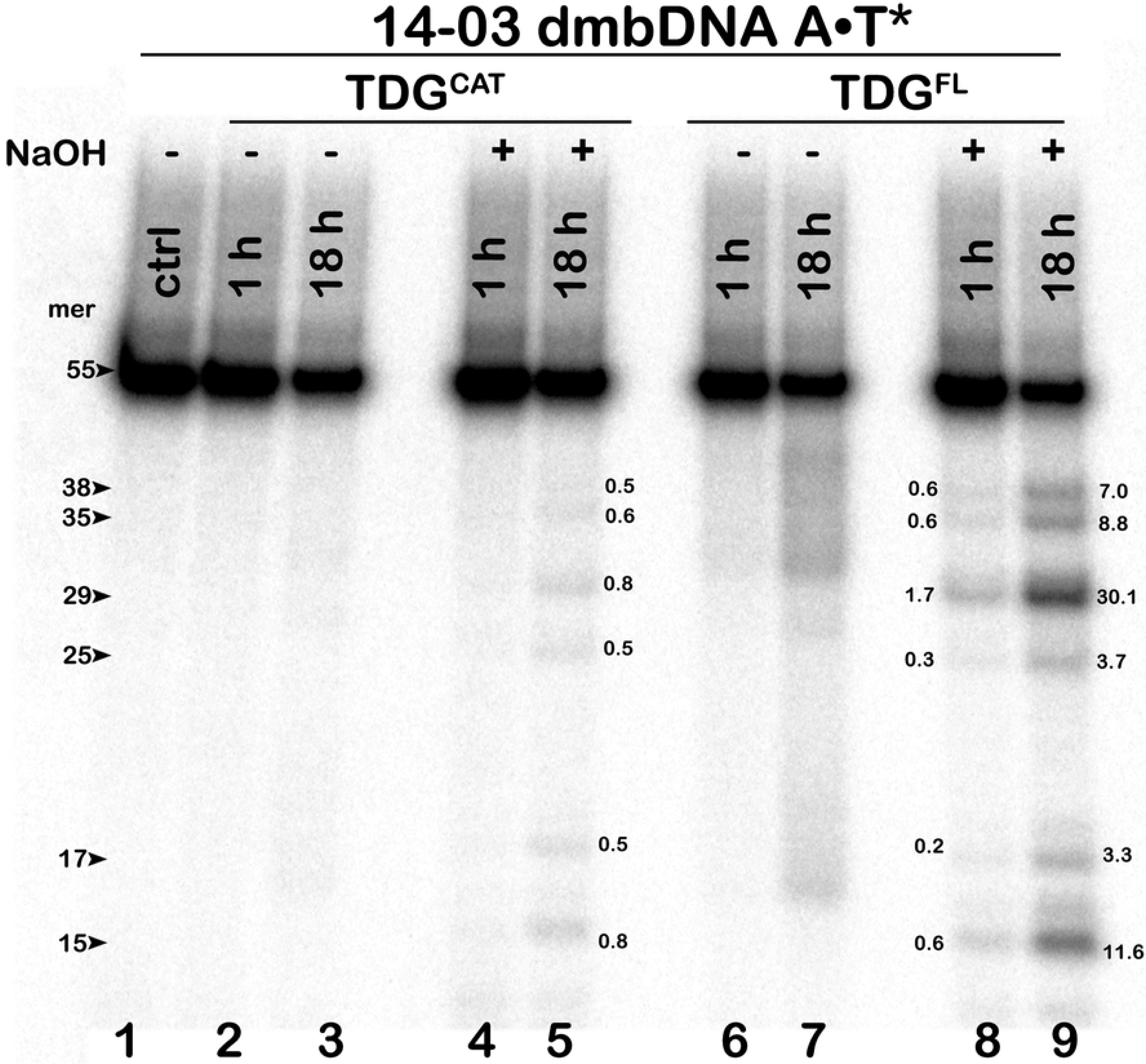
Denaturing PAGE analysis of the cleavage products following action of TDG^cat^ and TDG^FL^ on regular non-damaged 41 mer 14-03 A•T* dmbDNA substrate. After reaction samples were either treated or not by hot alkaline. Lane 1: control 14-03 dmbDNA with 5’-[^32^P]-labeled bottom 55 mer strand; lanes 2-3, same as 1, but with TDG^cat^ and without hot alkaline treatment; lanes 4-5, same as 2-3, but with hot alkaline treatment; lanes 6-9, same as 2-5, but with TDG^FL^. Arrows indicate the size of DNA substrate and cleavage fragments generated by TDG. The small numbers next to the bands in lanes 5, 8 and 9 correspond to the percentage of cleavage products. For details, see Materials and Methods.

n addition, we examined whether a functional analog of TDG, methyl-binding domain protein 4 (MBD4), exhibits any futile excision activity towards regular DNA duplexes. We incubated MBD4 with G•T* and A•T* duplexes for 1 h and 18 h, respectively. As expected, MBD4 efficiently excised mismatched T opposite to G but failed to exhibit detectable activity towards a regular DNA duplex (S1 Fig), suggesting the difference in the molecular mechanism of DNA damage recognition between these human G/T-specific DNA glycosylases.

### Kinetic parameters of the TDG^FL^-catalysed futile excision of T paired with A in a regular DNA duplex

We further characterized the substrate specificity of TDG^FL^ by comparing the kinetic parameters of cleavage of short blunt-ended 27-mer 14-03 and 24-mer 10-05 oligonucleotide duplexes containing either G•T* or A•T* base pair at defined positions in a TpG sequence context (the asterisk denotes either Cy5-labelled or 5′-^32^P-labelled DNA strand). Reactions were performed under single-turnover conditions using a 10-fold molar excess of the enzyme over the substrate, which provides the maximal rate of base excision (*k*obs) for a given substrate. As expected, the time course of the cleavage product accumulation showed that G•T is the preferred substrate for TDG^FL^; more than 80% of T at position 16 in the 14-03 G•T* duplex was excised in 5 min at 37°C (Fig. 3A, lanes 7 and 3B) in contrast to only 21% and 18% of T at two positions 16 and 12, respectively, in a regular 14-03 A•T* duplex after long incubation for 18 h at 37°C (Fig. 3C, lanes 9 and 3D). We also measured the kinetics of excision of T by TDG^FL^ in ^32^P-labelled 24-mer 10.05 G•T* and A•T* duplexes and observed similar large differences between the conventional and futile DNA glycosylase activities (S2 Fig). Based on these time courses, we measured *k*obs values of TDG-catalysed futile activity towards T in A•T base pair in duplex DNA and compared them to *k*obs of excision of T from a G•T mismatch using the same duplex. As shown in Table 2, when using the 14-03 duplex as a DNA substrate, the *k*obs values of TDG^FL^-catalysed excision of T opposite to regular A were, respectively, 330- and 270-fold lower than that of mismatched T in the 14.03 T•G duplex (0.47 min^−1^). When comparing the *k*obs values of TDG^FL^ on 10.05 DNA duplexes containing either G•T or A•T at the same position, the futile activity towards A•T base pair was 2.5–3-fold greater as compared to that in the 14-03 duplex, whereas the regular activity on a mismatched T in 10-05 was only 1.3-fold higher as compared to 14-03, resulting in only 85- and 100-fold differences between the G/T mismatch and futile activities in 10-05 duplexes (Table 2). These results suggest that the efficiency of TDG^FL^-catalysed futile excision as compared to the activities on the canonical DNA substrate strongly depends on the DNA sequence context.

**Figure 3.**
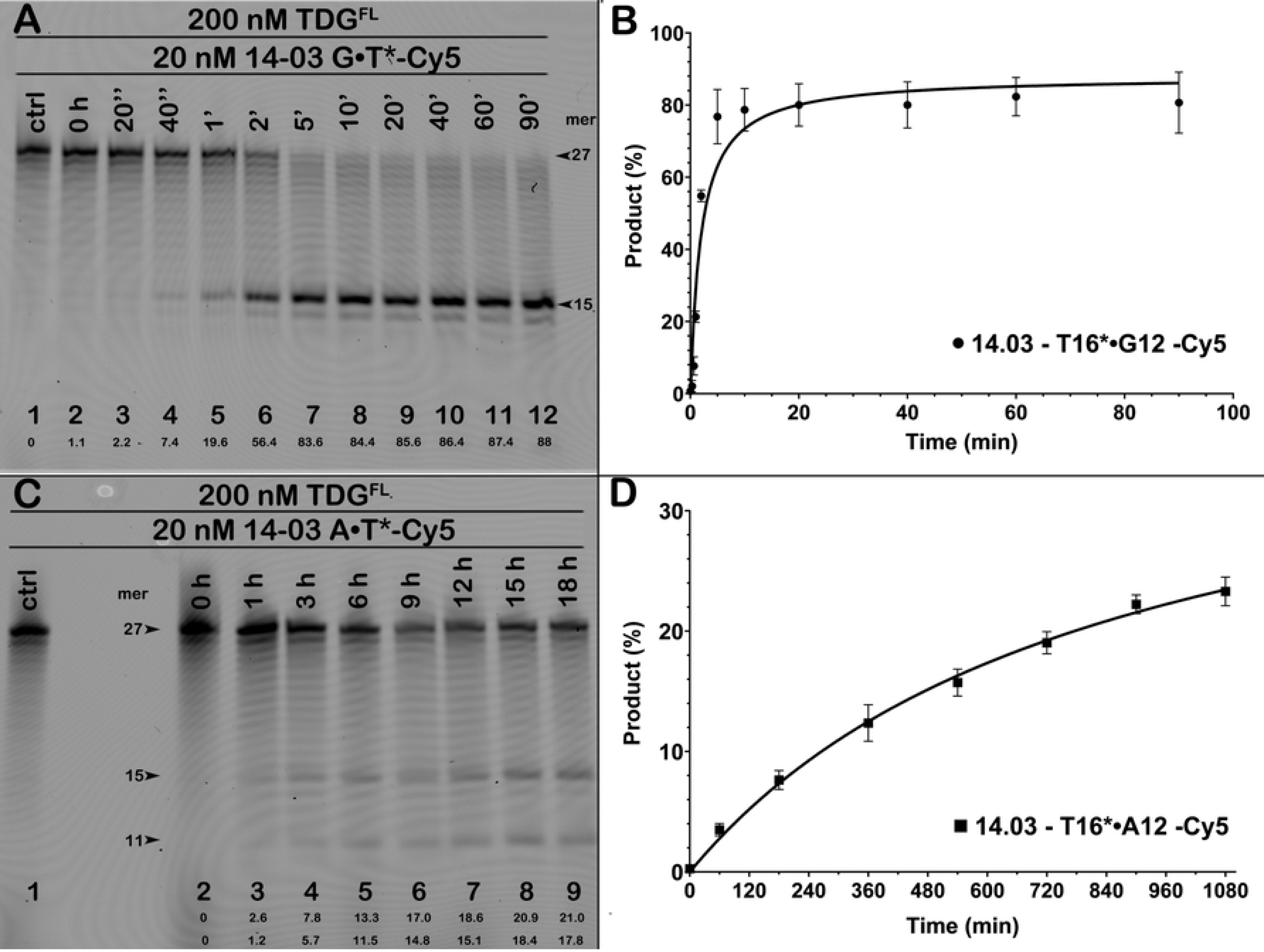
Time kinetics of TDG^FL^–catalysed excision of T from G•T and A•T base pairs in 27 mer 14-03 duplex oligonucleotide. (**A and C**) Denaturing PAGE analysis of the products of reaction. Time kinetics were performed using 200 nM TDG and 20 nM 5’-Cy5-labeled 27 mer 14-03 oligonucleotide duplex containing G•T or A•T base pairs at defined position. Arrows indicate the size of DNA substrate and cleavage fragments generated by TDG. The small numbers at bottom of the gel correspond to the percentage of cleavage products. (**B and D**) Graphic presentation of pre-steady-state single turnover kinetic of TDG^FL^-catalysed cleavage of 14-03 oligonucleotide duplexes. Each bar represents the mean values of TDG activity ± SD of three independent experiments. For details, see Materials and Methods.

Previously Morgan *et al.* demonstrated that TDG excises T from a 19-mer T•A duplex with an exceedingly low rate characterized by *k*max = 1.3×10^−5^ min^−1^, which was 17,600-fold slower than the activity on a T•G duplex (*k*max = 0.22 min^−1^) [28]. It should be noted that, in order to measure DNA glycosylase activity of TDG during long incubation time, the authors performed the reaction at 22°C to avoid rapid enzyme inactivation. In addition, they used denaturing HPLC at alkaline pH to separate the reaction products and measured their quantities by absorbance at 260 nm. Thus, the difference in the 100-fold preference for T•G vs T•A substrates observed in our work and the 10,000-fold preference observed in the previous studies might be explained by different temperature regimes. In a later study by Maiti *et al.* from the same laboratory, the futile activity of wild-type and mutant TDG proteins was measured using a ^32^P-labelled 28-mer T*•A duplex and denaturing PAGE to separate the reaction products [27]. In agreement with their previous study, these authors showed exceedingly weak activity of wild-type TDG towards a regular 28-mer T*•A duplex; after 4000 min at 22°C, the DNA glycosylase was able to excise only 3% of T residues paired with A. This led to a 20,000-fold difference between the *k*obs values of T excision measured on a 28-mer T•G *vs* T•A duplex [27]. It should be noted that the 28-mer duplex used in this study had a very particular sequence context with the cleaved strand highly enriched in purines with a single T in a TpG context.

To get insight into possible reasons for the observed discrepancies between our results and the published data [27, 28], we examined TDG^FL^-catalysed futile activity towards ^32^P-labelled 19- and 28-mer T*•A duplexes with DNA sequence contexts used in [27, 28] but employing our experimental conditions. After 18 h of incubation at 37°C, TDG^FL^ was able to excise 5.3 and 2.8% of T in T•A base pairs at positions 5 and 11, respectively, in the top strand of the 19-mer T*•A duplex and 9.9% of T at position 17 in the 28-mer T*•A duplex (S3 Fig). The measured efficiencies of TDG futile activity towards the regular 28-mer DNA duplex under our experimental conditions and that of Maiti *et al.* were very different: 9.9% of T excision in 1080 min (18 h) at 37°C *vs* 3% of T in 4000 min (66.7 h) at 22°C [27]. Based on these results, we suggest that the use of full-length version of the enzyme, DNA substrates with varying sequence contexts, physiologically relevant reaction temperatures and long incubation time play a critical role in the detection and characterization of the futile activity of human TDG.

### Long-term thermal stability of TDG

Time courses shown in Figure 3 and S2 Figure indicate that, even after 6, 9 and 12 h of incubation at 37°C, TDG^FL^ continues to excise pyrimidines in a regular DNA duplex, suggesting that the enzyme somehow maintains its activity despite the problem of long-term stability common for many enzymes under these conditions. Native TDG contains disordered N- and C-terminal tails, which may promote protein aggregation and loss of enzyme activity upon long incubation at 37°C [26]. To examine whether the intrinsically unfolded tails have an impact on the stability of TDG, we incubated TDG^cat^ and TDG^FL^ in the reaction buffer at 37°C for an extended time (0 h to 48 h) before adding a ^32^P-labelled 24-mer 10-13 T*•G duplex (a conventional TDG substrate) and incubating for another 1 h at 37°C to measure the remaining activitiy. As shown in Figure 4A-B, both truncated and native variants of TDG lost 90% of their activity after 12 h at 37°C (Fig. 4A, lanes 4 and 11). Noteworthy, TDG^FL^ exhibited somewhat higher stability with 13% and 3% remaining activity after 3 h and 12 h, respectively, while TDG^cat^ retained only 9.7% and 2.0% of its activity (lanes 10–11 *vs* lanes 3–4), suggesting slightly higher thermal stability of the native enzyme. Previously, Maiti *et al.* demonstrated that incubation of TDG in the presence of non-specific DNA prevents protein inactivation at 37°C for 2 h [26]. Therefore, we verified whether the presence of regular DNA duplex would affect the enzymes’ catalytic proficiency at 37°C. Incubation of 200 nM TDG proteins at 37°C in the presence of 20 nM unlabelled 24-mer 10-13 duplex slightly increased the stability of both variants (Fig. 4C, lanes 3–7 and 10–14, also Fig. 4D) as compared to the incubation without DNA (Fig. 4A, lanes 3–7 and 10–14, also Fig. 4B). Incubation of 200 nM TDG^cat^ with 200 nM 10-13 duplex did not affect its stability, as the activity of the truncated variant quickly dropped after 3 h of incubation and was completely lost after 24 h at 37°C (Fig. 4E, lanes 3 and 5). Unexpectedly, incubation of 200 nM TDG^FL^ at 37°C in the presence of an equimolar concentration of unlabelled 10-13 regular DNA duplex dramatically increased the enzyme stability (Fig. 4E, lanes 9–14, also Fig. 4F). TDG^FL^ exhibited full activity after 3 h at 37°C and was able to excise 12% of mismatched T even after 48-h incubation (Fig. 4E, lanes 10 and 13).

**Figure 4.**
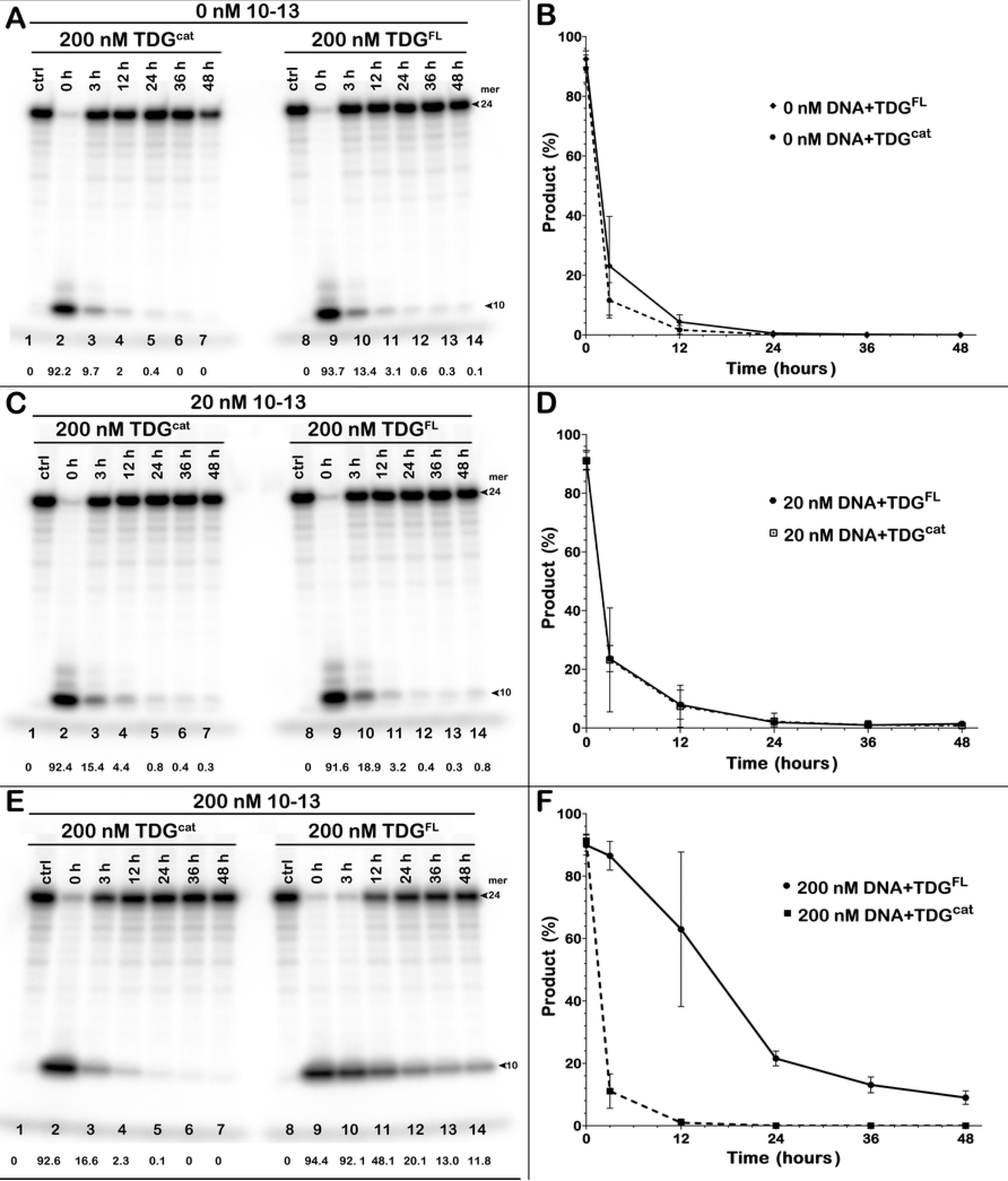
Impact of prolonged incubation of the TDG^cat^ and TDG^FL^ proteins at 37°C on their G/T mismatch specific DNA glycosylase activity. The proteins were incubated in the reaction buffer at 37°C for 0-48h with or without cold non-specific 24 mer 10-13 DNA duplex before adding 5’-[^32^P]-labelled 24 mer 10-05 G•T* duplex oligonucleotide and incubating further for 1h at 37°C. (**A, C, E**) Analysis of the TDG-catalysed cleavage products of 24 mer 10-05 G•T* duplex by denaturing PAGE. Before reaction with 10-05, 200 nM TDG was incubated at 37° in the presence of (**A**) 0 nM cold DNA; (**C**) 20 nM cold DNA; (**E**) 200 nM cold DNA. The small numbers at bottom of the gel correspond to the percentage of cleavage products. Arrows indicate substrate and cleavage product. (**B, D, F**) Graphic presentation of the remaining G/T-DNA glycosylase activity of the TDG^cat^ and TDG^FL^ proteins in the presence or absence of non-specific cold 24 mer 10-13 DNA duplex. Each bar represents the mean values of TDG activity ± SD of three independent experiments. For details, see Materials and Methods.

Overall, these results suggest that the presence of an equimolar amount of DNA duplex preserves the catalytic proficiency of a significant fraction of native TDG but not TDG^cat^ in the reaction. We propose that the presence of regular DNA stabilizes the protein with its N- and C-terminal tails, which may bind DNA duplex to prevent TDG aggregation and ensure its long-term stability at normal temperature of the human body. By contrast, TDG^cat^ lacks disordered tails and thus may be prone to self-aggregation at 37°C.

### Single-stranded oligoribonucleotides can modulate the futile activity of TDG

Recently, it has been shown that TDG binds G-rich single-stranded RNA with high affinity [49]. As a result, short and long RNA molecules with specific sequences can inhibit TDG-catalysed excision from U•G and T•G mismatches in duplex DNA [49]. In the present work, we examined the effect of short RNA oligonucleotides with varying sequences, taken from the previous study, on TDG-catalysed repair of 24-mer 10-05 T*•G and T*•A duplex oligonucleotides. As expected, the presence of RNAs that do not form stable complexes with TDG, such as 0.5–1.0 µM double-stranded G-rich RNA hairpin (GC-RNA) or single-stranded A-rich RNA (A30-RNA) [49], had little or no effect on the G/T-specific TDG activity on a 24-mer 10-05 T*•G duplex (Fig. 5A, lane 2 *vs* lanes 3–4 and 7–8; also Fig. 5B). In agreement with the published data [49], the presence of 0.5–1.0 µM modified G-rich RNA hairpin (Mut-RNA) containing an internal loop and a 3′-single-stranded tail, significantly inhibited TDG activity on the T*•G duplex down from 95% DNA cleavage in the control to 68–74% in the presence of RNA (Fig. 5A, lane 2 *vs* lanes 5–6; also Fig. 5B). As shown in Figure 5C, the presence of 0.5–1.0 µM high-affinity Mut-RNA strongly inhibited futile excision of T in a 24-mer 10-05 T*•A duplex down from 54% in the control to 40% and 12% in the presence of 0.5 µM and 1 µM RNA, respectively (Fig. 5C, lane 2 *vs* lanes 5–6). Mut-RNA inhibited the futile activity of TDG more strongly than its activity on a T•G mismatch, perhaps because of the differences between TDG binding affinities for RNA *vs* DNA. Comparing TDG binding constants for a T•G mismatch (*K*d = 18 nM), a T•A duplex (290 nM), Mut-RNA (140 nM), and GC-RNA (2900 nM) [49, 50], one can see that the affinity of TDG for Mut-RNA is 8-fold lower than for the T•G duplex but 21-fold higher than for the T•A duplex; therefore at the same concentration Mut-RNA will more strongly compete with TDG futile activity rather than with its G/T mismatch activity. Interestingly, unlike Mut-RNA, low-affinity GC-RNA and A30-RNA moderately stimulate TDG futile activity, up from 54% cleavage of a 24-mer 10-05 T*•A duplex in the control to 67–71% in the presence of RNA (Fig. 5C, lane 2 *vs* lanes 3–4 and 7– 8). These results suggest that non-specific, low-affinity binding of native TDG to dsRNA and G-poor ssRNA may stabilize the protein conformation and thus increase its thermal stability at 37°C, similar to what was observed with non-specific DNA duplexes. As expected, the presence of an equimolar concentration of unlabelled non-specific DNA did not influence the futile activity of TDG in a statistically significant manner (Fig. 5D).

**Figure 5.**
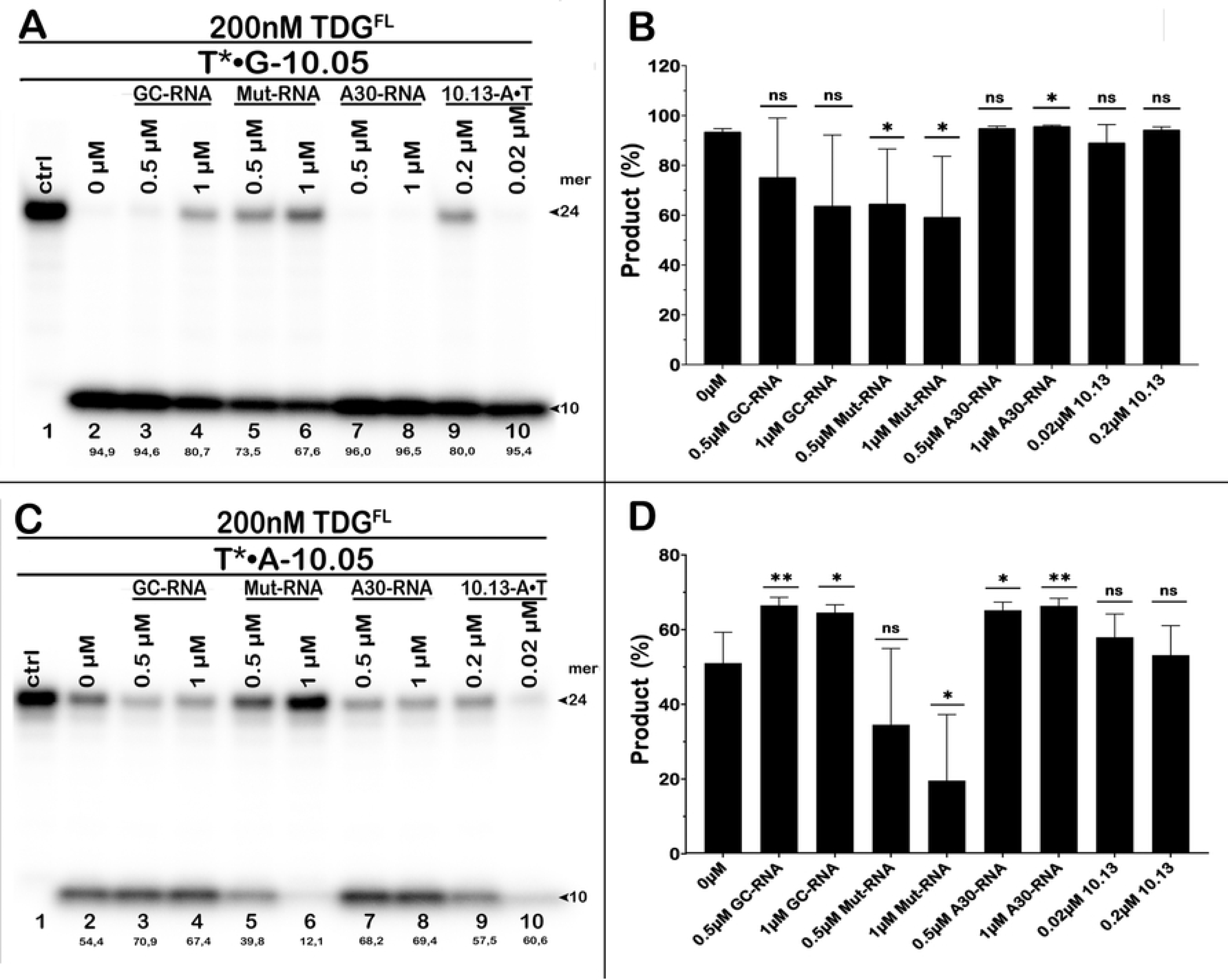
Impact of RNA oligonucleotides on G/T mismatch specific and futile DNA glycosylase activities of the TDG^FL^ protein. 24 mer 10-05 G•T* and A•T* duplex oligonucleotides were incubated with TDG^FL^ either in the presence of 0.5-1.0 µM RNA oligonucleotides or cold nonspecific 20-200 nM 10-13 DNA duplex at 37°C for 1h and 18h. (**A** and **C**) Analysis of the TDG-catalysed cleavage products of 24 mer 10-05 G•T* and A•T* duplex oligonucleotides, respectively, by denaturing PAGE. The small numbers at bottom of the gel correspond to the percentage of cleavage products. (**B** and **D**) Graphic presentation of the quantification of cleavage products shown on denaturing gels. Statistical significance of the differences was evaluated using two-tailed Student’s t-test (ns, not significant; ***, p < 0.001; **, p < 0.01; *, p < 0.05). For details, see Materials and Methods.

### The presence of G•T and G•U mispairs in DNA inhibit TDG-catalysed futile excision of neighbouring pyrimidines

Previously, it has been demonstrated that during TET/TDG-mediated active DNA demethylation at 5mCpG/5mCpG sites, DNA glycosylase-catalysed excision of 5caC is inhibited after base release and generation of an AP site in one of DNA strands [51]. Therefore, we decided to investigate whether the pattern of futile action is affected in the presence of T•G and U•G base pairs in the middle of a long 63-mer oligonucleotide duplex.

As shown in Figure 6, 18-h incubation of ^32^P-labelled regular 63-mer T*•A and T•A* duplexes with TDG^FL^ generates multiple cleavage fragments (55-, 44-, 38-, 20-, 16-, 13- and 7-mers in the T*•A and 57-, 54-, 45-, 29-, 25-, 23-, 10- and 7-mers in the T•A* substrate) which are due to the excision of pyrimidines (T and C) from TpG, CpA, CpG and TpA contexts (Fig. 6, lanes 2 and 4). As expected, incubation of a 63-mer T*•G duplex with TDG^FL^ generated a major 31-mer cleavage band indicating the excision of T at position 32 opposite to G (Fig. 6, lane 6). It should be noted that with the 5′-labeled T*•G substrate, longer futile excision products such as 55- and 44-mer fragments almost disappeared after 18 h while shorter ones (20-, 16-, 13- and 7-mer products) were still present (Fig. 6, lane 6 *vs* lane 2). The disappearance of 55- and 44-mer cleavage products (sites of pyrimidine excision located 3′ to the mismatch) is due to the very efficient excision of T at position 32, which makes 55- and 44-mer fragments invisible for detection by phosphoroimaging. Noteworthy, the generation of short cleavage fragments after incubation of TDG^FL^ with a 63-mer T*•G duplex suggests that tight binding of TDG^FL^ to the AP site product of excision of T opposite to G does not inhibit the futile activity at distances of 11 and more nucleotides 5′ of the mismatched T (Fig. 6, lane 6 *vs* lane 2). Nevertheless, the pattern of futile excision of pyrimidines by TDG^FL^ in the complementary DNA strand of the 5′-[^32^P]-labelled 63-mer T•G* duplex as compared to a regular T•A* duplex reveals a strong decrease in the generation of the 23-, 25-, 29- and 45-mer, but not of the 7-, 10-, 54- and 57-mer cleavage fragments (Fig. 6, lane 8 *vs* lane 4). These observations suggest strong inhibition at distances of 14 and 17 nucleotides 3′ of the mismatched G, and 2, 6 and 8 nucleotides 5′ of the mismatched G, when compared to the patterns of cleavage in a regular T•A* duplex (Fig. 6, lane 8 *vs* lane 4). Nevertheless, the futile activity of TDG^FL^ is not suppressed at longer distances from the mismatch: no inhibition was observed at 11 nucleotides (20-mer cleavage fragment) 5′ of the mismatched T in the top strand of the T*•G duplex and at 23 nucleotides (54-mer fragment) 3′ of the mismatched G and at 21 nucleotides (10-mer fragment) 5′ of the mismatched G in the bottom strand of T•G* duplex.

**Figure 6.**
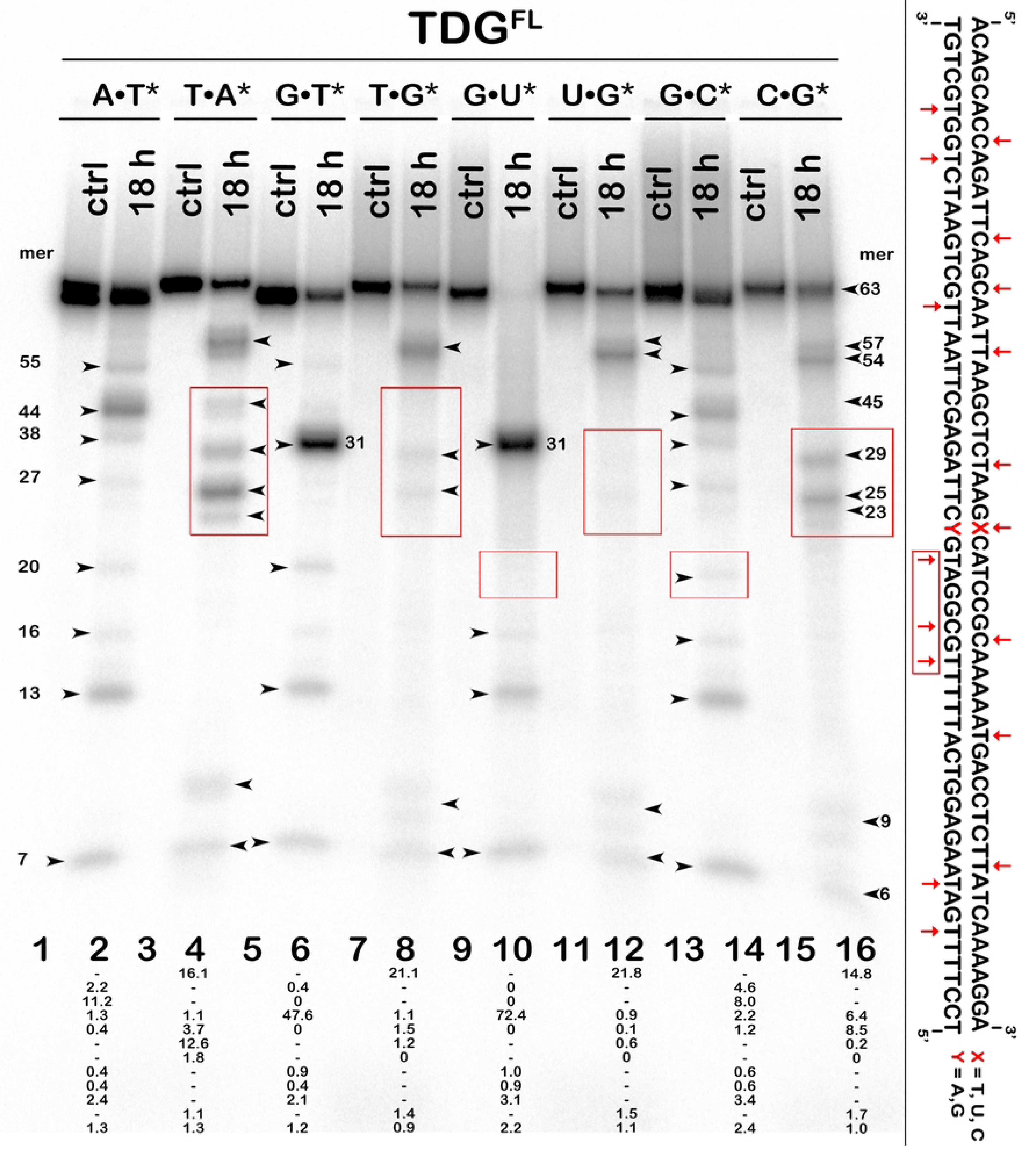
Action of TDG^FL^ on 63 mer A*•T, A•T* G•T*, T•G*, G•U*, U*•G, G•C*, C•G* duplex oligonucleotides, in which either top or bottom DNA strand is [^32^P]-labelled. Denaturing PAGE analysis of the cleavage products. The DNA bands framed in red boxes denote the difference in cleavage patterns between regular duplexes *vs* T•G and U•G duplexes. Arrows indicate the size of DNA substrate and cleavage fragments produced by TDG-catalysed base excision. The small numbers at bottom of the gel correspond to the percentage of cleavage products. The right panel shows schematic representation of the DNA sequence of 63 mer duplex oligonucleotide with red arrows depicting pyrimidines excised by TDG^FL^. For details, see Materials and Methods.

In addition, we compared TDG^FL^ catalysed excision patterns using 5′-[^32^P]-labelled 63-mer U*•G, U•G*, C*•G and C•G* duplexes (Fig. 6, lanes 10, 12, 14 and 16). TDG^FL^ excises mismatched U more efficiently than T (Fig. 6, lane 10 *vs* lane 6), resulting in a stronger inhibition of the futile activity towards neighbouring pyrimidine bases in a 63-mer U•G duplex when comparing to a regular C•G duplex (Fig. 6, lane 10 *vs* lane 14 and lane 12 *vs* lane 16). Because of the upstream truncation of the 63-mer due to the cleavage of U at position 32 in U*•G duplex, all downstream cleavage fragments (54, 44 and 38 nucleotides long), observed with a C*•G duplex, disappear completely (Fig. 6, lane 10 *vs* lane 14). Moreover, some upstream products of futile excision such as the 27- and 20-mer cleavage fragments also disappeared completely with the U*•G duplex (Fig. 6, lane 10 *vs* lane 14), suggesting strong inhibition of the futile activity on the U-containing strand at distances of 4 and 11 nucleotides 5′ of the uracil in the 63-mer. In addition, the products of futile excision 29, 25 and 23 nucleotides long, observed in the C•G* duplex, disappeared completely in the U•G* duplex (Fig. 6, lane 12 *vs* lane 16), showing strong inhibition of the futile activity at distances of 2, 6 and 8 nucleotides 5′ of the mismatched G in the bottom strand of the 63-mer DNA duplex. On the contrary, the presence of a U•G mispair did not suppress futile excision at longer distances, e. g., at 15 nucleotides 5′ of Ul (16-mer cleavage fragment) in the top strand of the U*•G duplex, and at 23 nucleotides 3′ of the mismatched G (54-mer fragment) and 21 nucleotides (10 mer cleavage fragment) 5′ of the mismatched G in the bottom strand of the U•G* duplex.

Taken together, these results suggest that the formation of a tight complex between TDG^FL^ and DNA duplex with an AP site following highly efficient excision of T and U opposite to G, strongly inhibits the TDG^FL^-catalysed futile activity io both strands at short distances (8–11 nucleotides) both 3′ and 5′ of T•G and U•G mismatches in duplex DNA.

### Futile activity of TDG towards enhancer sequences containing C, 5mC and 5hmC residues in a CpG context

Recently, it has been demonstrated that specific sets of enhancers in post-mitotic neurons are hotspots for single-strand breaks [45]. Therefore, we hypothesized that TDG could generate single-strand breaks in enhancers via its futile activity. To examine this, we measured TDG^FL^ activity towards enhancer DNA sequences such as neuronal enhancer element mm876 [52] and retinoic acid response element (RARE) involved in the regulation of the *Hic1* (Hypermethylated in Cancer 1) gene [53]. As expected, we observed a significant level of futile excision of T and C in TpG, CpA and CpG contexts in both DNA strands of a 63-mer mm876 duplex and a 27-mer RARE duplex (S4 and S5 Figs). These results may suggest that TDG would introduce SSBs in enhancers *in vivo* if it would be stably bound to DNA for a sufficiently long time.

TETs/TDG mediated removal of 5mC is known to be required for C/EBPα-induced pre-B-cell-to-macrophage differentiation [46]. Hence, we investigated whether TDG futile repair activity can excise the products of cytosine methylation and demethylation, 5mC and 5hmC, from two 27-mer oligonucleotide duplexes, RARE and 14.CpG (an artificial sequence context rich in CpG). Toward this end, 5′-[^32^P]-labelled 27-mer oligonucleotide duplexes RARE and 14.CpG containing C, 5hmC, or 5mC at position 15, were incubated with TDG^FL^ at 37°C for 1 h and 18 h. As shown in Figure 7, long-time 18 h incubation of unmodified RARE C*•G duplex generated 14- and 11-mer cleavage fragments indicating excision of 9.8% and 9.8% of C residues at positions 15 and 12, respectively (Fig. 7, lane 4). When TDG^FL^ was incubated with RARE 5hmC*•G and 5mC*•G duplexes for 18 h, the enzyme excised 18% of 5hmC and only 0.8% of 5mC residues at position 15 (Fig. 7, lanes 7 and 10); in other words, the presence of 5hmC in the RARE duplex stimulated TDG-catalysed futile excision 1.8-fold, whereas 5mC inhibited it more than 12-fold of C. Similar patterns of excision of C (20%), 5mC (1%) and 5hmC (27%) residues by TDG^FL^ were observed when using 14.CpG duplex (S6 Fig). Combined, these observations suggest that TDG may generate SSBs via the BER pathway in postmitotic cells in the enhancer DNA sequences containing 5hmC residues even without their conversion to 5fC or 5caC. Intriguingly, DNA methylation not only prevents transcription factors binding to enhancers but also strongly inhibits the DNA glycosylase-initiated futile excision of cytosines, thus avoiding appearance of persistent SSBs. We also surmise that in postmitotic cells, TETs-mediated generation of 5hmC in enhancer and promoter regions is sufficient for slow replication-independent DNA demethylation and generation of programmed DNA strand breaks via TDG-initiated BER.

**Figure 7.**
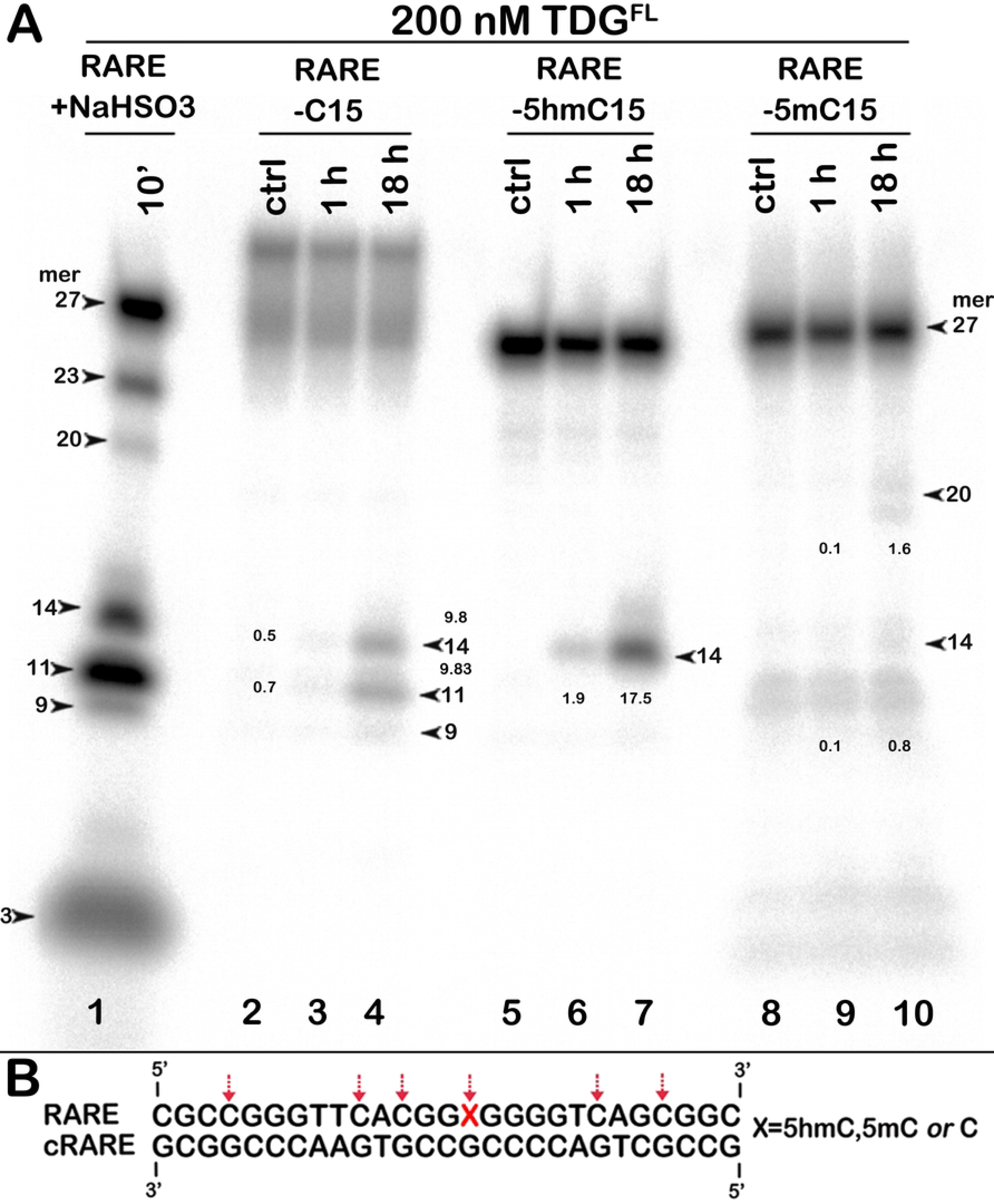
Action of TDG^FL^ towards 27 mer RARE duplex oligonucleotide containing C, 5hmC and 5mC at position 15. (**A**) Denaturing PAGE analysis of the cleavage products after incubation of TDG^FL^ with 5’-[^32^P]-labelled 27 mer RARE C*•G, 5hmC*•G and 5mC*•G duplex oligonucleotides for 1 and 18h at 37°C. Bisulfite-treated single-stranded 27 mer RARE oligonucleotide was incubated with hUNG to generate size markers denoting the cytosine positions. The small numbers next to the bands in lanes 3, 4, 6, 7, 9 and 10 correspond to the percentage of cleavage products. For details, see Materials and Methods. (**B**) Schematic representation of the DNA sequence of duplex oligonucleotide with arrows depicting pyrimidines excised by TDG^FL^.

### Bioinformatic analysis and structural modeling of full-length TDG

Intrigued by the ability of supposedly disordered N- and C-terminal tails of TDG to entail stabilization on the protein upon interaction with DNA, we explored possible reasons for this phenomenon. To figure out if there might be any communication between the N- and C-terminal tails of TDG, we have analysed the coevolution of amino acid properties in these regions. Correlated changes in physicochemical properties of residues residing far apart in the sequence often reflect functional interactions or dynamic physical contacts between them, which may be not evident from a static structure [54, 55]. Taking all TDG homologs from chordates and discarding those lacking the tails, we have used the CRASP tool [56] to search for correlations in the isoelectric point, local flexibility, hydrophobic index (Fig. 8A–C) and several other parameters reflecting hydrophobicity/hydrophilicity, chain flexibility, surface accessibility and tendency to form secondary structure (S7 Fig). As expected, no coevolution could be observed within the TDG catalytic core due to a high level of conservation and a small number of amino acid changes. The C-terminal tail was also almost free of either internal or cross-domain correlations. This is perhaps not surprising since the C-terminal tail provides a wide-area interface for sumoylation, which regulates many of TDG activities [57–59], and thus might coevolve instead with SUMO proteins or SUMO E3 ligases. However, multiple coevolving residue pairs were observed within the N-terminal tail, which also often showed correlations with residues at the junction of TDG^cat^ and the C-terminal tail (Fig. 8A–C and S7 Fig). This suggests that the N-terminal tail might be at least partially ordered and involved in some structural and/or functional cross-talk with other TDG regions.

**Figure 8.**
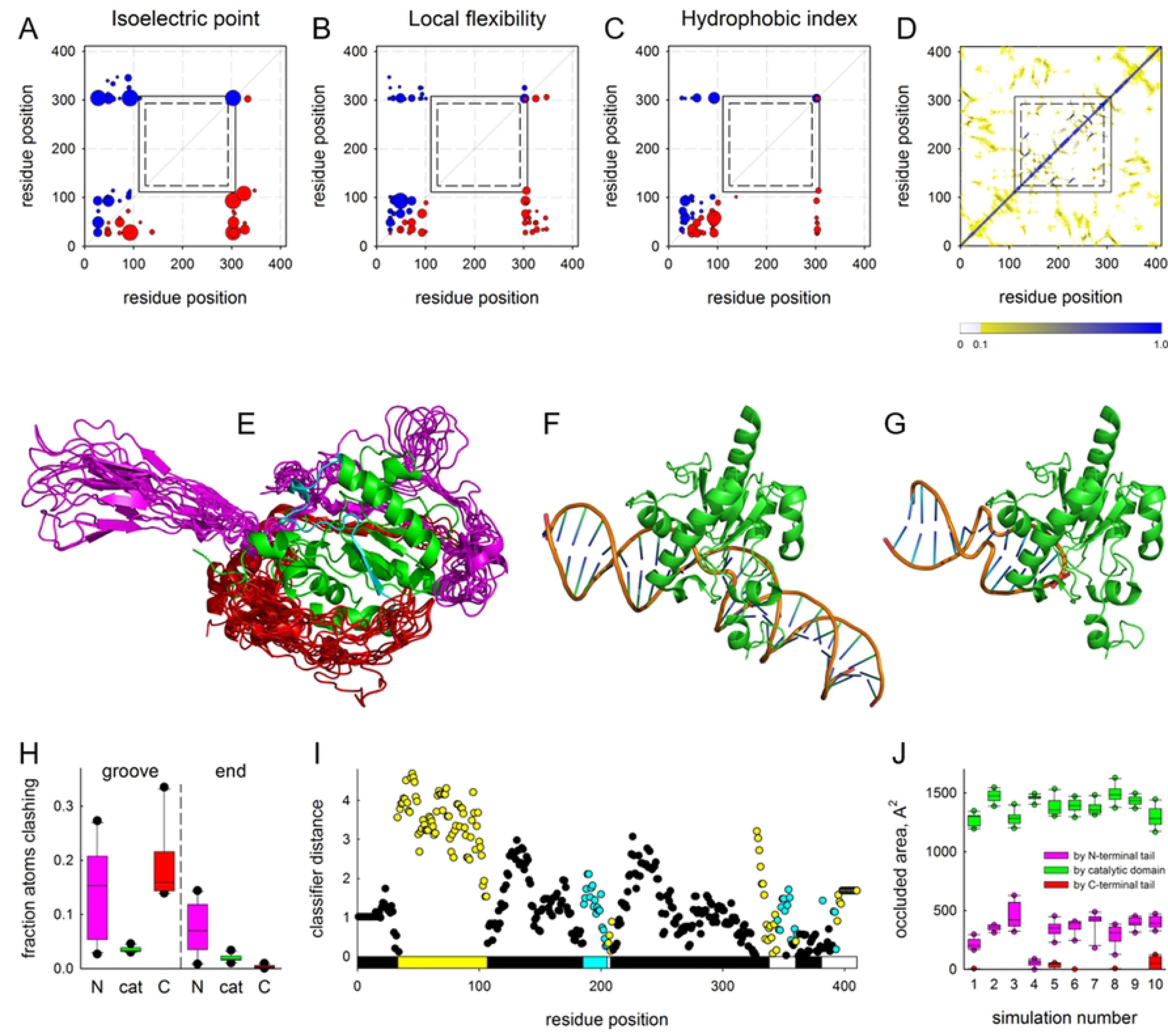
Analysis of structure and properties of TDG disordered tails. (**A–C**), correlation between the isoelectric point [75], local flexibility [76], and hydrophobic index [77] of TDG residues. Red circles under the diagonal, positive correlation; blue circles above the diagonal, negative correlation; circle radii are proportional to the absolute value of correlation coefficient (*r*ij) for the property for the given position pair. Only the pairs with *p* < 0.0001 (*r*ij > 0.1947) are shown. The rectangles delimit the catalytic domain [57] (residues 111–308; solid lines) and the UDG-F2_TDG_MUG conserved domain as defined in the NCBI Conserved Domain Database [78] (residues 124–293; dashed lines). (**D**), color-coded contact frequency map of TDG residues averaged over all coarse-grained simulations. (**E**), ten centroid structures from one coarse-grained simulation illustrating conformational variability of the TDG tails. The TDG^cat^ domain is shown in green as a single experimental structure (PDB ID 5HF7 [60]); the possible phase separation-prone fragment (residues 186– 205) is colored cyan. The cores of the simulated structures are superimposed over TDG^cat^ but omitted from the representation for clarity, leaving only the N-terminal tail (magenta) and the C-terminal tail (red). (**F, G**), positions of TDG^cat^ bound to the DNA minor groove (**F**) and to the duplex end (**G**); the latter is obtained by superimposing TDG^cat^ over the structure of Mug bound to the DNA end (PDB ID 1MTL, chain A [63]. (**H**), box plots showing the steric clashes between DNA and the N-terminal tail (N, magenta), TDG^cat^ (cat, green) and C-terminal tail (C, red) of the simulated structures docked in the groove and end positions. Clashes were counted as occurrences of a protein atom within 2.5 Å of any DNA atom. (**I**), ParSe predictions for TDG: cyan, residues intrinsically disordered and prone to undergo phase separation; yellow, intrinsically disordered but do not undergo phase separation; black, may or may not be intrinsically disordered but can fold to a stable conformation; white, no prediction. Bar shows TDG fragments with disorder predictions assigned based on the presence of 20 or more contiguous residues that are at least 90% of the same type. (**J**) area of the 186–205 fragment occluded by the N-terminal tail (magenta), TDG^cat^ (green) and C-terminal tail (red) of the model centroid structures in individual simulations.

At the next step, we sought to probe the conformational space available to full-length TDG. To do this, we started with an AlphaFold-generated model, in which both tails are present as extended chains with low confidence score (S8A Fig). The catalytic core is modeled by AlphaFold with a good confidence, with r.m.s.d. = 0.393 over the backbone atoms from the X-ray structure (PDB ID 5HF7) [60]. Since regular full-atom molecular dynamics is poorly suited to sample the huge number of conformations available to long unfolded protein chains, we resorted to coarse-grained Monte Carlo simulations. We used CABS-flex [61, 62], a modeling tool that implements a knowledge-based force field with parameters optimized to provide maximal convergence with molecular dynamics of proteins in explicit water at 300 K. Ten independent simulations with random seeds were run, and 1000 models were extracted and clustered for each one. Expectedly, TDG^cat^ was quite stable during the simulation (r.m.s.d. from the starting model 2.3–3.3 Å) whereas the tails coalesced from the fully unfolded state into better defined structures, which still demonstrated much higher conformational freedom than the core (Fig. 8D,E, and S8B-K Fig). Interestingly, antiparallel hairpin-like structures were often observed in the tails, especially the N-terminal one (Fig. 8D,E, and S8B–K Fig). Superimposing the models over the structure of TDG^cat^ bound to substrate DNA in the minor groove (Fig. 8F) [60] clearly reveals that a substantial conformational rearrangement of both tails would be required to avoid clashing with DNA (Fig. 8H). It is known, however, that Mug, the *E. coli* homolog of TDG lacking the tails, can bind to the end of a DNA duplex in a non-productive mode [63]. When we docked the models to the position of Mug occupying the DNA end (Fig. 8G), it significantly relieved clashes by the N-terminal tail (*p* < 0.005, Student’s paired *t*-test) and completely abolished them for the C-terminal tail (Fig. 8H). In the former, the atoms escaping from the clash were now more frequently found at distances between 2.5 Å and 4 Å from DNA (*p* < 0.05, Student’s paired *t*-test), which is more permissive for binding interactions. Also, the hairpin-like structures in the N-terminal tail in many cases could be fit into DNA grooves with some conformational adjustment.

Some, but not all intrinsically disordered protein regions are prone to undergo liquid–liquid phase separation, which is crucial for spatial organization of many processes, including DNA repair, inside the cell [64, 65]. TDG was reported to induce phase separation and condensation in chromatin both *in vitro* and in cells’ nuclei [66, 67]. To see whether the disorder in the TDG structure may assist phase separation, we used ParSe, a family of physics-based classifiers trained on a set of intrinsically disordered proteins to distinguish between condensate-forming and non-separating polypeptide regions [68]. While both tails were predicted to contain disorder-promoting residues of both types, only the N-terminal tail carried a contiguous stretch of such residues long enough to be classified as intrinsically disordered but not prone to phase separation (Fig. 8I and S9 Fig). Interestingly, both the basic ParSe algorithm and two extended classifiers with more physicochemical parameters considered robustly identified a fragment of TDG^cat^ (residues 186–205) as intrinsically disordered and prone to undergo phase separation. This region of the core was among the least stable in the simulations (r.m.s.f. plots in S8B–K Fig) and, even more intriguingly, is partly disordered in the structure of SUMO3-conjugated free TDG [59]. In the models, the 186–205 fragment contacts the N-terminal tail but not the C-terminal one (Fig. 8J), raising the possibility of induced disorder that may trigger a conformational change in TDG^cat^. Moreover, this fragment also interacts with the end of the duplex in the Mug-like binding mode.

The modeling and bioinformatic exercises allow us to propose a mechanism of TDG^FL^ protection by non-specific DNA. We suggest that TDG^FL^ binds ends of a DNA duplex (or, perhaps, even single-stranded DNA or RNA). In the free state, the conformationally relaxed, disorder-prone N-terminal tail might eventually induce disorder in the 186–205 fragment, possibly followed by TDG condensation. The core of isolated TDG^cat^ where the 186–205 fragment is directly exposed to solution, might also be prone to disorder. However, in the unproductive binding mode, the N-terminal tail is partially diverted to interactions with DNA whereas the 186–205 fragment is protected by its contacts with the duplex end. Thus, while the observed TDG^FL^ protection by non-specific DNA is clearly an *in vitro* phenomenon, it could reflect physiologically pertinent features of TDG related to order/disorder transitions in this important epigenetic regulator and guardian.

## Discussion

Over the course of evolution, DNA repair systems acquired high specificity for damaged DNA. Nevertheless, challenges to faithful discrimination of lesions among many orders of magnitude excess of regular bases in genomic DNA do exist. Under certain conditions, DNA repair can go astray and remove a regular DNA base; the same normal nucleotide is then incorporated, which may initiate multiple rounds of futile repair. Hence, futile base excision repair (f-BER) is useless if not mutagenic. In 1998, Seeberg and colleagues discovered that bacterial (AlkA), yeast (MAG) and human (ANPG) alkylpurine DNA glycosylases can excise regular adenine and guanine residues from nondamaged DNA duplex at measurable rates [40]. Increased spontaneous mutation rates were observed in *E. coli* overexpressing AlkA and MAG, respectively [40, 69]. In humans, an increased level of ANPG is linked with the risk of lung cancer, suggesting that futile BER may contribute to carcinogenesis [41]. Futile DNA repair activity was also shown for the bacterial and human nucleotide excision repair (NER), which can target regular DNA and excise nondamaged oligonucleotide fragments leading to futile excision/resynthesis cycles [70]. These futile excision activities of BER and NER are thought to act primarily on regular DNA, so their mutagenic effect in non-dividing cells may be limited by the error rates of repair DNA polymerases [71]. Intriguingly, when characterizing patterns of somatic mutations in post-mitotic neurons and polyclonal smooth muscle cells, it has been shown that neurons accumulate somatic mutations at a constant rate throughout life without cell division, with rates similar to mitotically active tissues [72, 73]. These observations suggest that mutations generated in the absence of DNA replication and cell division could be an important factor in somatic mutagenesis.

Here, we present evidence that native full-length TDG, TDG^FL^ can excise regular pyrimidines from T•A and C•G base pairs in duplex DNA in a futile manner with a significant efficiency (Figs 1 and 2). The rate constant of TDG^FL^-catalysed excision of T opposite to A are only 100–300-fold lower than that of mismatched T in a T•G duplex, suggesting that this futile activity could become physiologically significant upon long-time association of TDG with nucleosome-free DNA (Table 2). Notably, truncation of TDG down to its catalytic domain (TDG^cat^) lowers the activity on regular DNA duplexes more than tenfold, suggesting that the disordered N-(residues 1–110) and C-terminal (residues 309–410) tails play an important role in the futile activity. MBD4, a functional homolog of TDG, cannot excise pyrimidines from regular DNA, further substantiating a different mechanism of DNA substrate recognition by the latter DNA glycosylase (S1 Fig). Our data summarized in Table 2 are at odds with previous reports from Drohat’s laboratory, where excision of T from T•A base pairs and of C, 5mC and 5hmC opposite to G by TDG was measured at 22–23°C for 1 h to 50 h and was more than 10,000-fold slower than the excision of T from G•T mispairs [12, 24, 27, 28]. The reason for using the ambient temperature was that TDG irreversibly loses its activity at 37°C while remaining stable for at least 5 h at 22°C [26]. The drawback of using the low temperature is that the G/T-mismatch specific activity of TDG strongly depends on the temperature and is threefold higher at 37°C than at 22°C [26], suggesting that the futile activity of TDG becomes significant only at a physiologically relevant temperature.

**Table 2.** Pre-steady-state kinetic parameters of TDG-catalysed excision of T opposite to A or G in regular DNA duplexes.

| Substrate<sup>a</sup> | $k_{\text{obs}}$ (min<sup>-1</sup>) | $k_{\text{obs[G]}}/k_{\text{obs[A]}}$ |
| --- | --- | --- |
| 14-03 T16/G12, G•T* (TpG/CpG) | **0.47 ± 0.09** | **1** |
| 14-03 T12/A16, A•T* (TpG/CpA) | **0.0014 ± 0.0006** | **330** |
| 14-03 T16/A12, A•T* (TpG/CpA) | **0.0018 ± 0.0010** | **270** |
| 10-05 G14/T11, G•T* (TpG/CpG) | **0.61 ± 0.16** | **1** |
| 10-05 A14/T11, A•T* (TpG/CpA) | **0.0060 ± 0.0041** | **100** |
| 10-05 T11/G14, T*•G (TpG/CpG) | **0.59 ± 0.14** | **1** |
| 10-05 T11/A14, T*•A (TpG/CpA) | **0.0069 ± 0.0029** | **85** |

<sup>a</sup>Letters within the parentheses represent the sequence context.

Time courses of TDG^FL^-catalysed futile excision of pyrimidines (Fig. 3C,D) suggested to us that the native enzyme remains active even after 18 h of incubation at 37°C. Consistent with the previous work that showed stabilization of TDG in the presence of non-specific DNA [26], we found that the catalytic proficiency of TDG^FL^ is greatly stabilized by an equimolar amount of non-specific regular DNA duplex (Fig. 4). The native enzyme kept 100% and 10% of its G/T-specific DNA glycosylase activity after 3 h and 48 h at 37°C, respectively (Fig. 4E,F), At the same time, TDG^cat^ exhibited a drastic decrease of its G/T-specific activity after 3 h and completely lost it after 24 h at 37°C, and its stability was not influenced by non-specific DNA. Thus, the dramatic increase in the stability of native TDG in presence of DNA may explain in part the phenomenon of futile DNA repair. Seeking an explanation to this stabilization, we have also analysed the interactions and disorder of TDG tails using several bioinformatic and molecular modelling approaches. The models suggested that the structure of TDG is highly dynamic, with the N- and C-terminal tails sampling wide conformational space and undergoing rearrangements upon productive DNA binding. The N-terminal tail is presumably engaged in functional cross-talk with the catalytic domain, which could be important for phase separation involving TDG. Unproductive binding to DNA ends likely uncouples the N-terminal tail from TDG^cat^, stabilizing the full-length protein. It is tempting to speculate that MBD4 and other human DNA glycosylases, except for ANPG, do not exhibit detectable excision of regular DNA bases because of their limited conformational stability at 37°C. In addition, we showed that low-affinity small RNA oligonucleotides could stimulate TDG^FL^-catalysed futile excision via non-specific binding, which may stabilize protein conformation, whereas high-affinity RNA and DNA with T•G and U•G mismatches inhibits futile activity via strong binding and capturing the glycosylase by RNA and AP sites, respectively (Figs 5 and 6).

Recent study demonstrated that G/T-, but not G/U-specific activity of TDG is severely suppressed in DNA packaged into nucleosome core particles [74]. Therefore, our *in vitro* results may suggest that *in vivo* TDG could exhibit futile excision of pyrimidines in open chromatin regions (OCRs) depleted of nucleosomes in postmitotic non-dividing cells, which would result in the generation of persistent DNA strand breaks. Indeed, we show that *in vitro* TDG is able to excise, at a slow rate, cytosine residues in CpG-rich enhancers mm876 and RARE (S4 and S5 Figs). Intriguingly, recent studies showed that non-dividing neuronal cells accumulate high level of persistent SSBs in the enhancers at CpG dinucleotides associated with TET/TDG-mediated removal of 5mC [44, 45]. It has been suggested that cycles of DNA methylation coupled to active demethylation control the expression of genes involved in cell identity and can be the source of persistent SSBs in differentiated postmitotic neurons and macrophages [46]. Here, we demonstrate for the first time that native TDG excises 5hmC opposite to G in a CpG context with better efficiency as compared to regular C, whereas the presence of 5mC in duplex DNA strongly inhibits the futile activity (Fig. 7 and S6 Fig). Based on these results, we propose that *in vivo* TDG-catalysed BER pathway could participate in slow “active DNA demethylation” of genetic loci rich in 5hmC residues such as enhancers. 5hmC is a stable epigenetic mark typically enriched in transcriptionally active and tissue-specific genes and appears to be important for the maintenance of the cell differentiation status upon division [36–39]. It is tempting to speculate that TDG-mediated removal of 5hmC is part of epigenetic mechanisms, which regulate gene expression patterns and enable to preserve a specific cellular identity via generation of programmed SSBs.

## ABBREVIATIONS

TDG: human mismatch-specific thymine-DNA glycosylase
TDG^FL^: native, full-length version of TDG
TDG^cat^: truncated catalytic-domain TDG
BER: base excision repair
TET1-3: ten-eleven translocation family of proteins
MBD4: methyl-binding protein 4
hUNG: human uracil-DNA glycosylase
ANPG: human alkylpurine-DNA glycosylase
AP: apurinic/apyrimidinic
APE1: major human AP endonuclease 1
Xth: *E. coli* exonuclease III
Nfo: *E. coli* endonuclease IV
U: uracil
5mC: 5-methylcytosine
5hmC: 5-hydroxymethylcytosine
5fC: 5-formylcytosine
5caC: 5-carboxylcytosine
Hx: hypoxanthine
8oxoA: 7,8-dihydro-8-oxoadenine
3’-P: 3’-terminal phosphate
3’-PUA: 3’-terminal α,β-unsaturated phosphoaldehyde group
dmbDNA: dumbbell shaped duplex oligonucleotide.

## Author contributions

**Conceptualization:** D.M., S.T., D.O.Z., B.T.M., M.S.

**Funding acquisition:** S.T., B.T.M.., D.O.Z., M.S.

**Investigation:** D.M., B.D., D.O.Z., A.A.I., S.T.

**Methodology:** D.M., B.D., S.T., D.O.Z., A.A.I.

**Supervision:** S.T., B.T.M., M.S.

**Visualization:** D.M. B.D., S.T., D.O.Z., A.A.I., M.S.

**Writing ± original draft:** D.M., S.T., D.O.Z., M.S.

**Writing ± review & editing:** D.M., D.O.Z., M.S.

## ACKNOWLEDGEMENT

We would like to thank Dr Viktoryia Sidorenko and Arthur Grollman (State University of New York at Stony Brook, Stony Brook, U.S.A.) for their important role in initiating this work.

## Appendix A. Supporting Information

Supporting Information associated with this article can be found in the online version.

## References

1. Bird A. DNA methylation patterns and epigenetic memory. Genes Dev. 2002;16(1):6-21. Epub 2002/01/10. doi: 10.1101/gad.947102. PubMed PMID: 11782440.

2. Greenberg MVC, Bourc’his D. The diverse roles of DNA methylation in mammalian development and disease. Nat Rev Mol Cell Biol. 2019;20(10):590-607. Epub 2019/08/11. doi: 10.1038/s41580-019-0159-6. PubMed PMID: 31399642.

3. Smith ZD, Meissner A. DNA methylation: roles in mammalian development. Nat Rev Genet. 2013;14(3):204-20. Epub 2013/02/13. doi: 10.1038/nrg3354. PubMed PMID: 23400093.

4. Bird AP. DNA methylation and the frequency of CpG in animal DNA. Nucleic Acids Res. 1980;8(7):1499-504. PubMed PMID: 6253938.

5. Neddermann P, Jiricny J. The purification of a mismatch-specific thymine-DNA glycosylase from HeLa cells. J Biol Chem. 1993;268(28):21218-24. PubMed PMID: 8407958.

6. Hendrich B, Hardeland U, Ng HH, Jiricny J, Bird A. The thymine glycosylase MBD4 can bind to the product of deamination at methylated CpG sites. Nature. 1999;401(6750):301–4. PubMed PMID: 10499592.

7. Hitomi K, Iwai S, Tainer JA. The intricate structural chemistry of base excision repair machinery: implications for DNA damage recognition, removal, and repair. DNA Repair (Amst). 2007;6(4):410-28. PubMed PMID: 17208522.

8. Krokan HE, Bjoras M. Base excision repair. Cold Spring Harb Perspect Biol. 2013;5(4):a012583. Epub 2013/04/03. doi: 10.1101/cshperspect.a012583. PubMed PMID: 23545420; PubMed Central PMCID: PMC3683898.

9. Hang B, Medina M, Fraenkel-Conrat H, Singer B. A 55-kDa protein isolated from human cells shows DNA glycosylase activity toward 3,N4-ethenocytosine and the G/T mismatch. Proc Natl Acad Sci U S A. 1998;95(23):13561-6. PubMed PMID: 9811839.

10. Saparbaev M, Laval J. 3,N4-ethenocytosine, a highly mutagenic adduct, is a primary substrate for *Escherichia coli* double-stranded uracil-DNA glycosylase and human mismatch-specific thymine-DNA glycosylase. Proc Natl Acad Sci U S A. 1998;95(15):8508-13. PubMed PMID: 9671708.

11. Yoon JH, Iwai S, O’Connor TR, Pfeifer GP. Human thymine DNA glycosylase (TDG) and methyl-CpG-binding protein 4 (MBD4) excise thymine glycol (Tg) from a Tg:G mispair. Nucleic Acids Res. 2003;31(18):5399-404. PubMed PMID: 12954776.

12. Bennett MT, Rodgers MT, Hebert AS, Ruslander LE, Eisele L, Drohat AC. Specificity of human thymine DNA glycosylase depends on N-glycosidic bond stability. J Am Chem Soc. 2006;128(38):12510-9. PubMed PMID: 16984202.

13. Talhaoui I, Couve S, Ishchenko AA, Kunz C, Schar P, Saparbaev M. 7,8-Dihydro-8-oxoadenine, a highly mutagenic adduct, is repaired by Escherichia coli and human mismatch-specific uracil/thymine-DNA glycosylases. Nucleic Acids Res. 2013;41(2):912-23. Epub 2012/12/05. doi: 10.1093/nar/gks1149. PubMed PMID: 23209024; PubMed Central PMCID: PMC3553953.

14. Neddermann P, Jiricny J. Efficient removal of uracil from G.U mispairs by the mismatch-specific thymine DNA glycosylase from HeLa cells. Proc Natl Acad Sci U S A. 1994;91(5):1642-6. PubMed PMID: 8127859.

15. Hardeland U, Bentele M, Jiricny J, Schar P. The versatile thymine DNA-glycosylase: a comparative characterization of the human, Drosophila and fission yeast orthologs. Nucleic Acids Res. 2003;31(9):2261-71. Epub 2003/04/25. PubMed PMID: 12711670; PubMed Central PMCID: PMC154230.

16. Petronzelli F, Riccio A, Markham GD, Seeholzer SH, Genuardi M, Karbowski M, et al. Investigation of the substrate spectrum of the human mismatch-specific DNA N-glycosylase MED1 (MBD4): fundamental role of the catalytic domain. J Cell Physiol. 2000;185(3):473-80. PubMed PMID: 11056019.

17. Hashimoto H, Liu Y, Upadhyay AK, Chang Y, Howerton SB, Vertino PM, et al. Recognition and potential mechanisms for replication and erasure of cytosine hydroxymethylation. Nucleic Acids Res. 2012. Epub 2012/03/01. doi: 10.1093/nar/gks155. PubMed PMID: 22362737.

18. Cortazar D, Kunz C, Selfridge J, Lettieri T, Saito Y, MacDougall E, et al. Embryonic lethal phenotype reveals a function of TDG in maintaining epigenetic stability. Nature. 2011;470(7334):419–23. Epub 2011/02/01. doi: 10.1038/nature09672. PubMed PMID: 21278727.

19. Cortellino S, Xu J, Sannai M, Moore R, Caretti E, Cigliano A, et al. Thymine DNA glycosylase is essential for active DNA demethylation by linked deamination-base excision repair. Cell. 2011;146(1):67-79. Epub 2011/07/05. doi: 10.1016/j.cell.2011.06.020. PubMed PMID: 21722948; PubMed Central PMCID: PMC3230223.

20. Tahiliani M, Koh KP, Shen Y, Pastor WA, Bandukwala H, Brudno Y, et al. Conversion of 5-methylcytosine to 5-hydroxymethylcytosine in mammalian DNA by MLL partner TET1. Science. 2009;324(5929):930–5. Epub 2009/04/18. doi: 10.1126/science.1170116. PubMed PMID: 19372391; PubMed Central PMCID: PMC2715015.

21. Ito S, D’Alessio AC, Taranova OV, Hong K, Sowers LC, Zhang Y. Role of Tet proteins in 5mC to 5hmC conversion, ES-cell self-renewal and inner cell mass specification. Nature. 2010;466(7310):1129–33. Epub 2010/07/20. doi: 10.1038/nature09303. PubMed PMID: 20639862; PubMed Central PMCID: PMC3491567.

22. He YF, Li BZ, Li Z, Liu P, Wang Y, Tang Q, et al. Tet-mediated formation of 5-carboxylcytosine and its excision by TDG in mammalian DNA. Science. 2011;333(6047):1303–7. Epub 2011/08/06. doi: 10.1126/science.1210944. PubMed PMID: 21817016.

23. Ito S, Shen L, Dai Q, Wu SC, Collins LB, Swenberg JA, et al. Tet proteins can convert 5-methylcytosine to 5-formylcytosine and 5-carboxylcytosine. Science. 2011;333(6047):1300–3. Epub 2011/07/23. doi: 10.1126/science.1210597. PubMed PMID: 21778364; PubMed Central PMCID: PMC3495246.

24. Maiti A, Drohat AC. Thymine DNA glycosylase can rapidly excise 5-formylcytosine and 5-carboxylcytosine: potential implications for active demethylation of CpG sites. J Biol Chem. 2011;286(41):35334-8. Epub 2011/08/25. doi: 10.1074/jbc.C111.284620. PubMed PMID: 21862836; PubMed Central PMCID: PMC3195571.

25. Hardeland U, Kunz C, Focke F, Szadkowski M, Schar P. Cell cycle regulation as a mechanism for functional separation of the apparently redundant uracil DNA glycosylases TDG and UNG2. Nucleic Acids Res. 2007;35(11):3859-67. PubMed PMID: 17526518.

26. Maiti A, Drohat AC. Dependence of substrate binding and catalysis on pH, ionic strength, and temperature for thymine DNA glycosylase: Insights into recognition and processing of G.T mispairs. DNA Repair (Amst). 2011;10(5):545-53. Epub 2011/04/09. doi: 10.1016/j.dnarep.2011.03.004. PubMed PMID: 21474392; PubMed Central PMCID: PMC3084331.

27. Maiti A, Noon MS, MacKerell AD, Jr., Pozharski E, Drohat AC. Lesion processing by a repair enzyme is severely curtailed by residues needed to prevent aberrant activity on undamaged DNA. Proc Natl Acad Sci U S A. 2012;109(21):8091-6. Epub 2012/05/11. doi: 10.1073/pnas.1201010109. PubMed PMID: 22573813; PubMed Central PMCID: PMC3361372.

28. Morgan MT, Bennett MT, Drohat AC. Excision of 5-Halogenated Uracils by Human Thymine DNA Glycosylase: ROBUST ACTIVITY FOR DNA CONTEXTS OTHER THAN CpG. J Biol Chem. 2007;282(38):27578-86. PubMed PMID: 17602166.

29. Wu H, Wu X, Shen L, Zhang Y. Single-base resolution analysis of active DNA demethylation using methylase-assisted bisulfite sequencing. Nat Biotechnol. 2014;32(12):1231-40. Epub 2014/11/05. doi: 10.1038/nbt.3073. PubMed PMID: 25362244; PubMed Central PMCID: PMC4269366.

30. Wagner M, Steinbacher J, Kraus TF, Michalakis S, Hackner B, Pfaffeneder T, et al. Age-dependent levels of 5-methyl-, 5-hydroxymethyl-, and 5-formylcytosine in human and mouse brain tissues. Angew Chem Int Ed Engl. 2015;54(42):12511-4. Epub 2015/07/04. doi: 10.1002/anie.201502722. PubMed PMID: 26137924; PubMed Central PMCID: PMC4643189.

31. Caldwell BA, Liu MY, Prasasya RD, Wang T, DeNizio JE, Leu NA, et al. Functionally distinct roles for TET-oxidized 5-methylcytosine bases in somatic reprogramming to pluripotency. Mol Cell. 2021;81(4):859-69.e8. Epub 2020/12/23. doi: 10.1016/j.molcel.2020.11.045. PubMed PMID: 33352108; PubMed Central PMCID: PMCPMC7897302.

32. Bachman M, Uribe-Lewis S, Yang X, Williams M, Murrell A, Balasubramanian S. 5-Hydroxymethylcytosine is a predominantly stable DNA modification. Nat Chem. 2014;6(12):1049-55. Epub 2014/11/21. doi: 10.1038/nchem.2064. PubMed PMID: 25411882; PubMed Central PMCID: PMCPMC4382525.

33. Wei A, Zhang H, Qiu Q, Fabyanic EB, Hu P, Wu H. 5-hydroxymethylcytosines regulate gene expression as a passive DNA demethylation resisting epigenetic mark in proliferative somatic cells. bioRxiv. 2023. Epub 2023/10/09. doi: 10.1101/2023.09.26.559662. PubMed PMID: 37808741; PubMed Central PMCID: PMCPMC10557716.

34. Lian CG, Xu Y, Ceol C, Wu F, Larson A, Dresser K, et al. Loss of 5-hydroxymethylcytosine is an epigenetic hallmark of melanoma. Cell. 2012;150(6):1135-46. Epub 2012/09/18. doi: 10.1016/j.cell.2012.07.033. PubMed PMID: 22980977; PubMed Central PMCID: PMCPMC3770275.

35. Jin SG, Jiang Y, Qiu R, Rauch TA, Wang Y, Schackert G, et al. 5-Hydroxymethylcytosine is strongly depleted in human cancers but its levels do not correlate with IDH1 mutations. Cancer Res. 2011;71(24):7360-5. Epub 2011/11/05. doi: 10.1158/0008-5472.Can-11-2023. PubMed PMID: 22052461; PubMed Central PMCID: PMCPMC3242933.

36. Spruijt CG, Gnerlich F, Smits AH, Pfaffeneder T, Jansen PW, Bauer C, et al. Dynamic readers for 5-(hydroxy)methylcytosine and its oxidized derivatives. Cell. 2013;152(5):1146-59. Epub 2013/02/26. doi: 10.1016/j.cell.2013.02.004. PubMed PMID: 23434322.

37. Nestor CE, Ottaviano R, Reddington J, Sproul D, Reinhardt D, Dunican D, et al. Tissue type is a major modifier of the 5-hydroxymethylcytosine content of human genes. Genome Res. 2012;22(3):467-77. Epub 2011/11/23. doi: 10.1101/gr.126417.111. PubMed PMID: 22106369; PubMed Central PMCID: PMCPMC3290782.

38. Stroud H, Feng S, Morey Kinney S, Pradhan S, Jacobsen SE. 5-Hydroxymethylcytosine is associated with enhancers and gene bodies in human embryonic stem cells. Genome Biol. 2011;12(6):R54. Epub 2011/06/22. doi: 10.1186/gb-2011-12-6-r54. PubMed PMID: 21689397; PubMed Central PMCID: PMCPMC3218842.

39. Cui XL, Nie J, Ku J, Dougherty U, West-Szymanski DC, Collin F, et al. A human tissue map of 5-hydroxymethylcytosines exhibits tissue specificity through gene and enhancer modulation. Nat Commun. 2020;11(1):6161. Epub 2020/12/04. doi: 10.1038/s41467-020-20001-w. PubMed PMID: 33268789; PubMed Central PMCID: PMCPMC7710742.

40. Berdal KG, Johansen RF, Seeberg E. Release of normal bases from intact DNA by a native DNA repair enzyme. Embo J. 1998;17(2):363-7. PubMed PMID: 9430628.

41. Leitner-Dagan Y, Sevilya Z, Pinchev M, Kramer R, Elinger D, Roisman LC, et al. N-methylpurine DNA glycosylase and OGG1 DNA repair activities: opposite associations with lung cancer risk. J Natl Cancer Inst. 2012;104(22):1765-9. Epub 2012/10/30. doi: 10.1093/jnci/djs445. PubMed PMID: 23104324; PubMed Central PMCID: PMC3502197.

42. Talhaoui I, Couve S, Gros L, Ishchenko AA, Matkarimov B, Saparbaev MK. Aberrant repair initiated by mismatch-specific thymine-DNA glycosylases provides a mechanism for the mutational bias observed in CpG islands. Nucleic Acids Res. 2014;42(10):6300-13. Epub 2014/04/03. doi: 10.1093/nar/gku246. PubMed PMID: 24692658; PubMed Central PMCID: PMC4041421.

43. Puc J, Aggarwal AK, Rosenfeld MG. Physiological functions of programmed DNA breaks in signal-induced transcription. Nat Rev Mol Cell Biol. 2017;18(8):471-6. Epub 2017/05/26. doi: 10.1038/nrm.2017.43. PubMed PMID: 28537575.

44. Reid DA, Reed PJ, Schlachetzki JCM, Nitulescu, II, Chou G, Tsui EC, et al. Incorporation of a nucleoside analog maps genome repair sites in postmitotic human neurons. Science. 2021;372(6537):91–4. Epub 2021/04/03. doi: 10.1126/science.abb9032. PubMed PMID: 33795458; PubMed Central PMCID: PMCPMC9179101.

45. Wu W, Hill SE, Nathan WJ, Paiano J, Callen E, Wang D, et al. Neuronal enhancers are hotspots for DNA single-strand break repair. Nature. 2021;593(7859):440–4. Epub 2021/03/27. doi: 10.1038/s41586-021-03468-5. PubMed PMID: 33767446; PubMed Central PMCID: PMCPMC9827709.

46. Wang D, Wu W, Callen E, Pavani R, Zolnerowich N, Kodali S, et al. Active DNA demethylation promotes cell fate specification and the DNA damage response. Science. 2022;378(6623):983–9. Epub 2022/12/02. doi: 10.1126/science.add9838. PubMed PMID: 36454826; PubMed Central PMCID: PMCPMC10196940.

47. Morera S, Grin I, Vigouroux A, Couve S, Henriot V, Saparbaev M, et al. Biochemical and structural characterization of the glycosylase domain of MBD4 bound to thymine and 5-hydroxymethyuracil-containing DNA. Nucleic Acids Res. 2012;40(19):9917-26. Epub 2012/08/01. doi: 10.1093/nar/gks714. PubMed PMID: 22848106; PubMed Central PMCID: PMC3479182.

48. Gelin A, Redrejo-Rodriguez M, Laval J, Fedorova OS, Saparbaev M, Ishchenko AA. Genetic and biochemical characterization of human AP endonuclease 1 mutants deficient in nucleotide incision repair activity. PLoS One. 2010;5(8):e12241. Epub 2010/09/03. doi: 10.1371/journal.pone.0012241. PubMed PMID: 20808930; PubMed Central PMCID: PMC2923195.

49. McGregor LA, Zhu B, Goetz AM, Sczepanski JT. Thymine DNA glycosylase is an RNA-binding protein with high selectivity for G-rich sequences. J Biol Chem. 2023;299(4):104590. Epub 2023/03/09. doi: 10.1016/j.jbc.2023.104590. PubMed PMID: 36889585; PubMed Central PMCID: PMCPMC10124917.

50. Morgan MT, Maiti A, Fitzgerald ME, Drohat AC. Stoichiometry and affinity for thymine DNA glycosylase binding to specific and nonspecific DNA. Nucleic Acids Res. 2011;39(6):2319-29. Epub 2010/11/26. doi: 10.1093/nar/gkq1164. PubMed PMID: 21097883; PubMed Central PMCID: PMC3064789.

51. Weber AR, Krawczyk C, Robertson AB, Kuśnierczyk A, Vågbø CB, Schuermann D, et al. Biochemical reconstitution of TET1-TDG-BER-dependent active DNA demethylation reveals a highly coordinated mechanism. Nat Commun. 2016;7:10806. Epub 2016/03/05. doi: 10.1038/ncomms10806. PubMed PMID: 26932196; PubMed Central PMCID: PMCPMC4778062.

52. Song M, Yang X, Ren X, Maliskova L, Li B, Jones IR, et al. Mapping cis-regulatory chromatin contacts in neural cells links neuropsychiatric disorder risk variants to target genes. Nat Genet. 2019;51(8):1252-62. Epub 2019/08/02. doi: 10.1038/s41588-019-0472-1. PubMed PMID: 31367015; PubMed Central PMCID: PMCPMC6677164.

53. Hassan HM, Kolendowski B, Isovic M, Bose K, Dranse HJ, Sampaio AV, et al. Regulation of Active DNA Demethylation through RAR-Mediated Recruitment of a TET/TDG Complex. Cell Rep. 2017;19(8):1685-97. Epub 2017/05/26. doi: 10.1016/j.celrep.2017.05.007. PubMed PMID: 28538185.

54. Little DY, Chen L. Identification of coevolving residues and coevolution potentials emphasizing structure, bond formation and catalytic coordination in protein evolution. PLoS One. 2009;4(3):e4762. Epub 2009/03/11. doi: 10.1371/journal.pone.0004762. PubMed PMID: 19274093; PubMed Central PMCID: PMCPMC2651771.

55. Sutto L, Marsili S, Valencia A, Gervasio FL. From residue coevolution to protein conformational ensembles and functional dynamics. Proc Natl Acad Sci U S A. 2015;112(44):13567-72. Epub 2015/10/22. doi: 10.1073/pnas.1508584112. PubMed PMID: 26487681; PubMed Central PMCID: PMCPMC4640757.

56. Afonnikov DA, Kolchanov NA. CRASP: a program for analysis of coordinated substitutions in multiple alignments of protein sequences. Nucleic Acids Res. 2004;32(Web Server issue):W64-8. Epub 2004/06/25. doi: 10.1093/nar/gkh451. PubMed PMID: 15215352; PubMed Central PMCID: PMCPMC441589.

57. Steinacher R, Schar P. Functionality of human thymine DNA glycosylase requires SUMO-regulated changes in protein conformation. Curr Biol. 2005;15(7):616-23. Epub 2005/04/13. doi: 10.1016/j.cub.2005.02.054. PubMed PMID: 15823533.

58. Baba D, Maita N, Jee JG, Uchimura Y, Saitoh H, Sugasawa K, et al. Crystal structure of thymine DNA glycosylase conjugated to SUMO-1. Nature. 2005;435(7044):979–82. Epub 2005/06/17. doi: 10.1038/nature03634. PubMed PMID: 15959518.

59. Baba D, Maita N, Jee JG, Uchimura Y, Saitoh H, Sugasawa K, et al. Crystal structure of SUMO-3-modified thymine-DNA glycosylase. J Mol Biol. 2006;359(1):137-47. Epub 2006/04/22. doi: 10.1016/j.jmb.2006.03.036. PubMed PMID: 16626738.

60. Coey CT, Malik SS, Pidugu LS, Varney KM, Pozharski E, Drohat AC. Structural basis of damage recognition by thymine DNA glycosylase: Key roles for N-terminal residues. Nucleic Acids Res. 2016;44(21):10248-58. Epub 2016/09/02. doi: 10.1093/nar/gkw768. PubMed PMID: 27580719; PubMed Central PMCID: PMC5137436.

61. Jamroz M, Orozco M, Kolinski A, Kmiecik S. Consistent View of Protein Fluctuations from All-Atom Molecular Dynamics and Coarse-Grained Dynamics with Knowledge-Based Force-Field. J Chem Theory Comput. 2013;9(1):119-25. Epub 2013/01/08. doi: 10.1021/ct300854w. PubMed PMID: 26589015.

62. Kuriata A, Gierut AM, Oleniecki T, Ciemny MP, Kolinski A, Kurcinski M, et al. CABS-flex 2.0: a web server for fast simulations of flexibility of protein structures. Nucleic Acids Res. 2018;46(W1):W338-w43. Epub 2018/05/16. doi: 10.1093/nar/gky356. PubMed PMID: 29762700; PubMed Central PMCID: PMCPMC6031000.

63. Barrett TE, Savva R, Barlow T, Brown T, Jiricny J, Pearl LH. Structure of a DNA base-excision product resembling a cisplatin inter-strand adduct. Nat Struct Biol. 1998;5(8):697-701. PubMed PMID: 9699633.

64. Chong PA, Forman-Kay JD. Liquid-liquid phase separation in cellular signaling systems. Curr Opin Struct Biol. 2016;41:180-6. Epub 2016/08/24. doi: 10.1016/j.sbi.2016.08.001. PubMed PMID: 27552079.

65. Spegg V, Altmeyer M. Biomolecular condensates at sites of DNA damage: More than just a phase. DNA Repair (Amst). 2021;106:103179. Epub 2021/07/27. doi: 10.1016/j.dnarep.2021.103179. PubMed PMID: 34311273; PubMed Central PMCID: PMCPMC7612016.

66. Deckard CE, 3rd, Banerjee DR, Sczepanski JT. Chromatin Structure and the Pioneering Transcription Factor FOXA1 Regulate TDG-Mediated Removal of 5-Formylcytosine from DNA. J Am Chem Soc. 2019;141(36):14110-4. Epub 2019/08/29. doi: 10.1021/jacs.9b07576. PubMed PMID: 31460763.

67. McGregor LA, Deckard CE, 3rd, Smolen JA, Porter GM, Sczepanski JT. Thymine DNA glycosylase mediates chromatin phase separation in a DNA methylation-dependent manner. J Biol Chem. 2023;299(7):104907. Epub 2023/06/13. doi: 10.1016/j.jbc.2023.104907. PubMed PMID: 37307918; PubMed Central PMCID: PMCPMC10404674.

68. Ibrahim AY, Khaodeuanepheng NP, Amarasekara DL, Correia JJ, Lewis KA, Fitzkee NC, et al. Intrinsically disordered regions that drive phase separation form a robustly distinct protein class. J Biol Chem. 2023;299(1):102801. Epub 2022/12/18. doi: 10.1016/j.jbc.2022.102801. PubMed PMID: 36528065; PubMed Central PMCID: PMCPMC9860499.

69. Xiao W, Samson L. In vivo evidence for endogenous DNA alkylation damage as a source of spontaneous mutation in eukaryotic cells. Proc Natl Acad Sci U S A. 1993;90(6):2117-21. Epub 1993/03/15. PubMed PMID: 7681584; PubMed Central PMCID: PMC46036.

70. Branum ME, Reardon JT, Sancar A. DNA repair excision nuclease attacks undamaged DNA. A potential source of spontaneous mutations. J Biol Chem. 2001;276(27):25421-6. PubMed PMID: 11353769.

71. Chan KK, Zhang QM, Dianov GL. Base excision repair fidelity in normal and cancer cells. Mutagenesis. 2006;21(3):173-8. PubMed PMID: 16613912.

72. Lodato MA, Rodin RE, Bohrson CL, Coulter ME, Barton AR, Kwon M, et al. Aging and neurodegeneration are associated with increased mutations in single human neurons. Science. 2018;359(6375):555-9. Epub 2017/12/09. doi: 10.1126/science.aao4426. PubMed PMID: 29217584; PubMed Central PMCID: PMCPMC5831169.

73. Moore L, Cagan A, Coorens THH, Neville MDC, Sanghvi R, Sanders MA, et al. The mutational landscape of human somatic and germline cells. Nature. 2021;597(7876):381–6. Epub 2021/08/27. doi: 10.1038/s41586-021-03822-7. PubMed PMID: 34433962.

74. Tarantino ME, Dow BJ, Drohat AC, Delaney S. Nucleosomes and the three glycosylases: High, medium, and low levels of excision by the uracil DNA glycosylase superfamily. DNA Repair (Amst). 2018;72:56-63. Epub 2018/10/01. doi: 10.1016/j.dnarep.2018.09.008. PubMed PMID: 30268365; PubMed Central PMCID: PMCPMC6420825.

75. Zimmerman JM, Eliezer N, Simha R. The characterization of amino acid sequences in proteins by statistical methods. J Theor Biol. 1968;21(2):170-201. Epub 1968/11/01. doi: 10.1016/0022-5193(68)90069-6. PubMed PMID: 5700434.

76. Ragone R, Facchiano F, Facchiano A, Facchiano AM, Colonna G. Flexibility plot of proteins. Protein Eng. 1989;2(7):497-504. Epub 1989/05/01. doi: 10.1093/protein/2.7.497. PubMed PMID: 2748566.

77. Ponnuswamy PK, Prabhakaran M, Manavalan P. Hydrophobic packing and spatial arrangement of amino acid residues in globular proteins. Biochim Biophys Acta. 1980;623(2):301-16. Epub 1980/06/26. doi: 10.1016/0005-2795(80)90258-5. PubMed PMID: 7397216.

78. Marchler-Bauer A, Derbyshire MK, Gonzales NR, Lu S, Chitsaz F, Geer LY, et al. CDD: NCBI’s conserved domain database. Nucleic Acids Res. 2015;43(Database issue):D222-6. Epub 2014/11/22. doi: 10.1093/nar/gku1221. PubMed PMID: 25414356; PubMed Central PMCID: PMCPMC4383992.

